# Deep Linear Modeling of Hierarchical Functional Connectivity in the Human Brain

**DOI:** 10.1101/2020.12.13.422538

**Authors:** Wei Zhang, Eva Palacios, Pratik Mukherjee

## Abstract

The human brain exhibits hierarchical modular organization, which is not depicted by conventional fMRI functional connectivity reconstruction methods such as independent component analysis (ICA). To map hierarchical brain connectivity networks (BCNs), we propose a novel class of deep (multilayer) linear models that are constructed such that each successive layer decomposes the features of the preceding layer. Three of these are multilayer variants of Sparse Dictionary Learning (SDL), Non-Negative Matrix Factorization (NMF) and Fast ICA (FICA). We present a fourth deep linear model, Deep Matrix Fitting (MF), which incorporates both rank reduction for data-driven hyperparameter determination as well as a distributed optimization function. We also introduce a novel framework for theoretical comparison of these deep linear models based on their combination of mathematical operators, the predictions of which are tested using simulated resting state fMRI data with known ground truth BCNs. Consistent with the theoretical predictions, Deep MF and Deep SDL performed best for connectivity estimation of 1^st^ layer networks, whereas Deep FICA and Deep NMF were modestly better for spatial mapping. Deep MF provided the best overall performance, including computational speed. These deep linear models can efficiently map hierarchical BCNs without requiring the manual hyperparameter tuning, extensive fMRI training data or high-performance computing infrastructure needed by deep nonlinear models, such as convolutional neural networks (CNNs) or deep belief networks (DBNs), and their results are also more explainable from their mathematical structure. These benefits gain in importance as continual improvements in the spatial and temporal resolution of fMRI reveal more of the hierarchy of spatiotemporal brain architecture. These new models of hierarchical BCNs may also advance the development of fMRI diagnostic and prognostic biomarkers, given the recent recognition of disparities between low-level vs high-level network connectivity across a wide range of neurological and psychiatric disorders.

## 1. Introduction

Functional Magnetic Resonance Imaging (fMRI) has been widely used for the identification of brain connectivity networks (BCNs) (Bartels et al., 2005; Beckmann et al., 2005; Biswal et al., 1995, 2010; Bullmore et al., 2009; Duncan et al., 2010; Stam et al., 2014). A variety of scientific investigations have already demonstrated the hierarchical modular organization of human brain networks (Bassett et al., 2008; Biswal et al., 2005; Bullmore et al., 2009; Sporns et al., 2004). The architecture of cortical BCNs is organized at different spatial scales, from both functional and structural perspectives, ranging from local circuits at the microscale to columns and layers at the mesoscale to areas and areal networks at the macroscale (Bullmore et al., 2009; Power et al., 2011; Stam et al., 2014; Sporns et al., 2004).

In the last two decades, a variety of computational methods have been developed to detect BCNs, e.g., General Linear Modeling (GLM), Graph Theory, Independent Component Analysis (ICA), and Sparse Dictionary Learning (SDL) (Andersen et al., 1999; Calhoun et al., 2001; Lee et al., 2011; Lee et al., 2016; Lv et al., 2015; Zhang et al., 2017; Zhang et al., 2018; Zhang et al., 2019). However, these methods are based on a ‘shallow’ framework that cannot identify in unsupervised data-driven fashion the hierarchical and spatially overlapping organization of BCNs using resting-state fMRI (rsfMRI) or task-evoked fMRI (tfMRI) signals (Hu et al., 2018; Huang et al., 2018; Zhang et al., 2019; Zhang et al., 2020). Traditionally, the hierarchical spatial organization of BCNs has been indicated by varying the number of features in shallow linear models, for example from low to high numbers of independent components in ICA (Iraji et al., 2019; Smith et al., 2009), and noting that smaller networks at the more granular decomposition tend to merge or otherwise recombine to form larger networks at the coarser decomposition. However, there is no principled, unsupervised way to map this hierarchical organization with shallow methods.

Fortunately, with the advent of deep learning, algorithms have been developed that are capable of reconstructing hierarchical network architectures, e.g., the Deep Convolutional Auto Encoder (DCAE), Deep Belief Network (DBN) and Convolutional Neural Network (CNN) (Bengio et al., 2012; Esteva, et al., 2019; Gurovich et al., 2019; Hannun et al., 2019; LeCun et al., 2015; Plis et al., 2014; Schmidhuber et al., 2015; Suk et al., 2014; Suk et al., 2016; Zhang et al., 2020). The Restricted Boltzmann Machine (RBM) can be used to model fMRI time series signals and effectively reconstruct functional brain networks with impressive accuracy (Hu et al., 2018; Huang et al., 2018). Moreover, other recent studies reported meaningfully hierarchical temporal organization of tfMRI time series, each with corresponding task-evoked BCNs (Hu et al., 2018; Zhang et al., 2019; Zhang et al., 2020) using DCAE, RBM and DBN. In general, these machine learning techniques are considered to be deep nonlinear models, e.g., deep neural networks (DNN). Although these nonlinear models such as DBN have recently proven effective at hierarchical spatiotemporal decomposition of task-evoked fMRI data (Dong et al., 2020), there are several disadvantages: (i) large training samples; (ii) extensive computational resources, e.g., graphics processing units (GPUs) or tensor processing units (TPUs); (iii) manual tuning of hyperparameters; (iv) time-consuming training process; (v) non-convergence to the global optimum; and (vi) “black box” results that lack explainability. Deep linear algorithms can overcome all these shortcomings of nonlinear techniques, since they are fast even on conventional central processing units (CPUs) with hyperparameters that can be automatically determined and with convex optimization functions that are guaranteed to converge. Furthermore, as we show in the theoretical analysis below, important aspects of their behavior can be explained from their relatively simple mathematical structure. For fMRI research, these deep linear models can detect BCNs using data from relatively few experimental subjects compared to deep nonlinear models and may prove especially useful and efficient as the spatial and temporal resolution of fMRI continues to improve, revealing more of the hierarchy of brain organization.

For hierarchical spatial functional connectivity mapping, we adopt a compositional approach to develop multilayer versions of SDL (Deep SDL), Fast ICA (Deep FICA; Seo, 2018) and Non-negative Matrix Factorization (Deep NMF; Trigeorgis et al., 2016), as well as a novel multilayer linear model that we name Deep Matrix Fitting (Deep MF). We contrast these deep linear algorithms for fMRI functional connectivity analysis in two ways. First, we employ theory to investigate the mathematical properties of these models, in order to predict differences in, e.g., network sparsity, connectivity strength and convergence velocity. Second, we conduct *in silico* connectivity reconstruction experiments using simulated fMRI signal time series to test the predictions of the theoretical analyses for the relative performance of the four deep linear models for fMRI brain network mapping. This leads to clear conclusions about the strengths and weaknesses of each method and provides a guide to the research community for applying this novel class of network reconstruction methodologies as well as for developing new ones.

## 2. Methods

### 2.1 Shallow versus Deep Linear Models of fMRI Functional Connectivity

The following introductions in Sections 2.2 to 2.5 provide the fundamentals of each deep linear model. Furthermore, these descriptions are prerequisites to analyze the theoretical properties of each model in the succeeding sections. Figure 1 compares the computational steps of a deep linear model versus that of a conventional “shallow” linear model:

**Figure 1.**
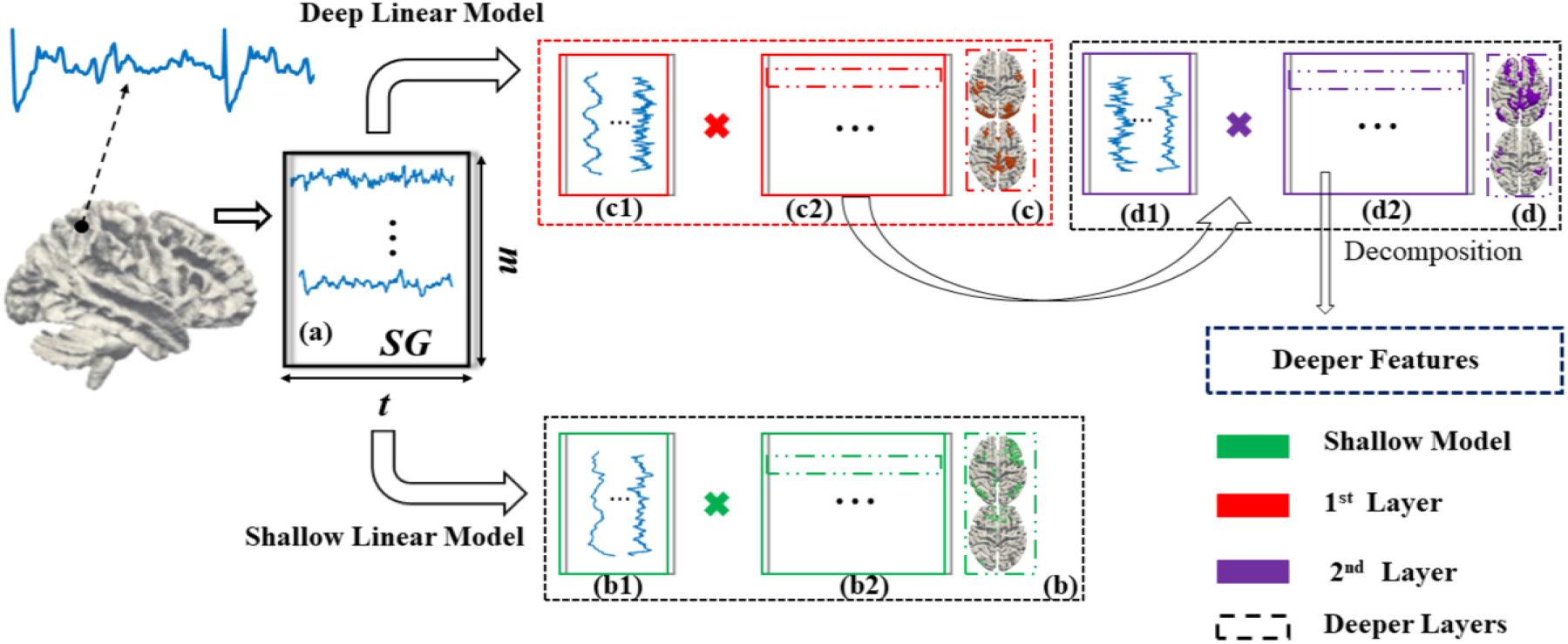
Deep linear model versus shallow linear model. The shallow model has only a single layer, i.e., a single decomposition. The deep linear model is constructed via multiple layers, i.e., continuous decomposition. (**a**) *SG* represents the input fMRI signal matrix; it contains the *t* time points and *m* voxels. (**b**) describes the pipeline of a shallow model in which the original input signal is decomposed into the weight matrix/dictionary (shown as **b1**) and feature matrix, i.e. connectivity networks (shown as **b2**). (**c**) and (**d**) represent the 1^st^ and 2^nd^ layers of the linear deep model, respectively. (**c1**) represents the 1st layer weight matrix/dictionary identified via SG. (**c2**) represents the 1^st^ layer feature matrix, i.e. connectivity networks, recognized via SG. Similarly, (**d1**) and (**d2**) represent the corresponding matrices of the 2^nd^ layer, which are both derived from the 1^st^ layer feature matrix. The dashed blue rectangle indicates the deeper features beyond the 2^nd^ layer that are derived from the 2^nd^ layer feature matrix.

### 2.2 Deep Matrix Fitting

We propose a novel and efficient deep linear model that we name Deep Matrix Fitting. This algorithm aims to detect the hierarchical and overlapping organization of BCNs better than previously described data-driven functional connectivity reconstruction methods, e.g., ICA, SDL, DCAE and DBN (Calhoun et al., 2001; Lv et al., 2015; Zhang et al., 2020; Hinton and Salakhutdinov, 2006; Hinton et al., 2012). Due to the constraints of spatial independence in ICA, some investigators have reported that ICA cannot easily identify extensively overlapped functional brain networks (Calhoun et al., 2001; McKeown and Sejnowski, 1998; Zhang et al., 2019). Although SDL can efficiently derive spatial features, i.e., functional brain networks, based on rsfMRI and tfMRI, it is very challenging to leverage the dictionary size, sparsity trade off and even number of layers to implement a deep SDL. To be specific, one must heuristically estimate the dictionary size and number of layers. Simply utilizing the same size of dictionary and number of layers can easily result in the vanishing of spatial features of deeper layers, due to iteratively using the ℓ_1_ norm. Recent deep nonlinear models, such as DBN, can successfully reveal the architecture of hierarchical spatiotemporal features. Unfortunately, the probabilistic energy-based model of DBNs necessarily requires a large number of training samples to avoid overfitting. Furthermore, DBN requires extensive computational resources such as GPUs and even TPUs (Zhang et al., 2019; Zhang et al., 2020). The novel Deep MF proposed in this work successfully solves these aforementioned problems. Deep MF can automatically estimate the optimal dictionary size, sparsity trade-off and number of layers, using an operator of rank reduction (Wen et al., 2012; Shen et al., 2014). In other words, Deep MF does not require any manual hyperparameter tuning to decompose the rsfMRI signal matrix. Since Deep MF is a deep linear model, it should detect latent features faster than DBN while only requiring conventional CPUs. In general, Deep MF can be approximately considered as a deep SDL (described in Section 2.3) with the additional mechanism to automatically determine all crucial hyperparameters via rank reduction.

The equation governing Deep MF is:

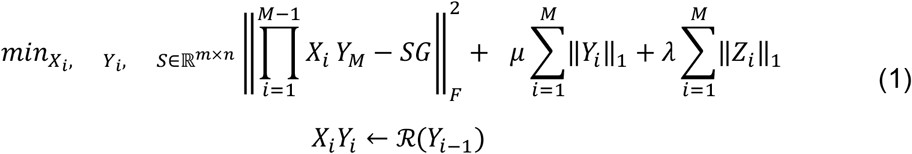

where 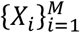 represents the hierarchical dictionaries, e.g., *X_i_* indicates the dictionary of the *i* th layer. 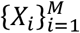 is also considered as the time series in GLM and the weight matrix in ICA and DBN. *M* is the total number of layers. Similarly, 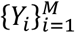 represents the hierarchical spatial features, e.g., *Y_i_* indicates the spatial features of *i^th^* layer. 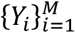 is also denoted as a correlation matrix. 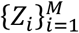 are the matrices of background components, which is usually treated as the noise. 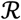 represents a rank reduction operator (RRO) to automatically estimate the hyperparameters and more details will be introduced in the following section. Naturally, we assume the spatial features *Y*_*i*−1_ can be decomposed as deeper dictionary *X_i_* and spatial features *Y_i_*, in order to implement the deep linear framework (Figure 1). Therefore, the original input data *SG* can be decomposed as 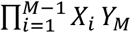. In Eq. (1), λ and *μ* are known as the sparse trade off to control the sparsity levels of background components and spatial features, respectively. In addition, in Eq. (1), ‖∙‖_*F*_ and ‖∙‖_1_ represent the Frobenius and ℓ_1_ norms, respectively.

This optimization function, shown as Eq. (1), consists of more parameters than ICA and SDL. In general, SDL includes two parameters to be optimized: dictionary and correlation matrix. Naturally, it is easier to comprehensively employ alternative optimizer and shrinkage methods (Wen et al., 2012). Before optimizing Eq. (1), we need to convert Eq. (1) to an augmented Lagrangian function. If considering the *k*^th^ layer, we have

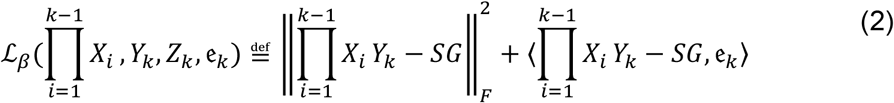

For Eq. (2), for *k* layers (we assume the total number of layers as *k*), these can be solved using Alternating Direction of Method of Multipliers (ADMM) (Shen et al., 2014), and to solve 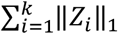, we jointly utilize the shrinkage method. In Eq. (2), all parameters are as discussed before, with 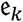 defined as the multiplier. The ℓ_1_ norm of *Y_k_* and *Z_k_* shown in Eq. (1) can be solved directly using the shrinkage method (Beck et al., 2009).

The iterative format to solve Eq. (2) using ADMM can be organized as follows:

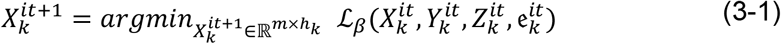

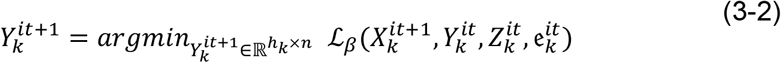

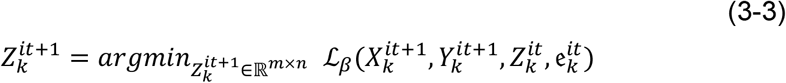

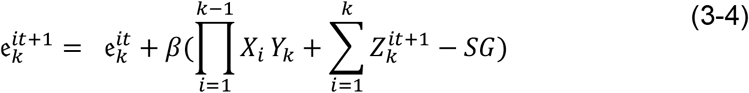

Using ADMM, in each iteration (the current iteration is represented as *it*), we update a single parameter independently, and finally calculate the multiplier, based on the current error. Since, in each single step, only one parameter is optimized and others are fixed, Eq (2) is considered as a convex problem and the global optimum can be obtained via a descent algorithm, e.g. gradient descent (GD) or ADMM. In Eq. (3-4), *β* denotes the step length.

To automatically estimate the dictionary size and number of layers, we introduce the operator RRO. Briefly, RRO focuses on the identification of major components included in the raw data, and simultaneously determines which components are relatively weak and that will therefore be continuously merged into the background matrices. In general, RRO demonstrates that the number of units, i.e., dictionary size, should be consistently reduced, if considering deeper layers (Hinton et al., 2012; Zhang et al., 2019). In other words, the continuous increase of units in deeper layers can result in lack of convergence. If the number of units/dictionary size, i.e., the estimated rank of the matrix, is reduced to one, that indicates the decomposition should be terminated. Hence, the layer that owns a rank of unity should be considered the final layer. Deep MF employs RRO to continuously reduce the dictionary size and therefore also determine the number of layers. In fact, it does not require any manual design for the essential hyperparameters of deep learning models, such as the number of layers or unit number of each layer that are used in DBN and other peer deep models.

In detail, this rank estimator RRO employs a technique of rank-revealing by continuously using orthogonal decomposition, in this case via *QR* factorization (Wen et al., 2012; Shen et al., 2014). The advantage of *QR* is that it is faster and makes fewer requirements of the input matrix. For example, *QR* performs orthogonal decomposition faster than Singular Value Decomposition (*SVD*) and can solve incomplete (i.e., number of features < number of samples) and over-complete (i.e., number of features > number of samples) matrices.

At the beginning, *r*\* is denoted as the initial estimated rank of *S^i^* and we denote *r* as the optimal rank estimation of input matrix *S^i^*. If *r*\*≥*r* holds, the detection of the diagonal line of the upper-triangular matrix in the *QR* factorization can be performed using the input matrix *S^i^*. If we can determine the ideal size of *QR* factorization using *S^i^* in the work with permutation matrix *E*, the diagonal matrix *R* is non-increasing in magnitude (Wen et al., 2012; Shen et al., 2014). The *QR* factorization and rank-revealing will eventually provide a reasonable solution using a proper thresholding value introduced in Eq. (2) and Eq. (3) (Wen et al., 2012; Shen et al., 2014). By detecting the diagonal line of matrix *R*, we compute two vectors *d* ∈ ℝ^*r*^ and *r* ∈ ℝ^*r*−1^:

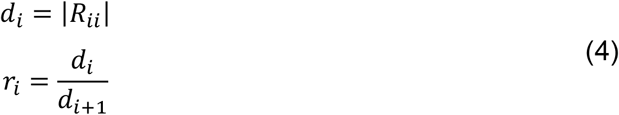

And then examine the value:

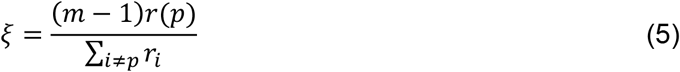

where *r(p)* is the maximum element of the vector *r* (with the largest index *p* if the maximum value is not unique). In our current implementation, we reset the rank estimated *r*, if *ξ* > 2, and this adjustment can be successfully done only once (Wen et al., 2012; Shen et al., 2014).

The mathematical definition of RRO is shown below:

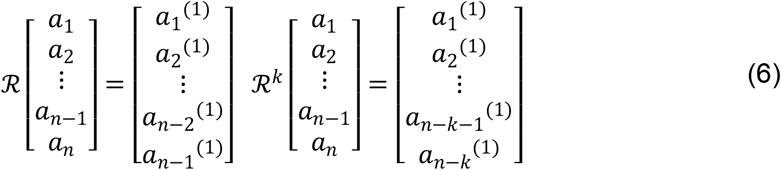

where 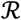 denotes the RRO operator; and theoretically, we have 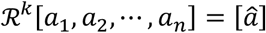, if *k* → ∞. It clearly demonstrates that the RRO can continuously maintain the vital components and reduce the dimensions of the original data. By continuously using the technique of low rank estimation, Deep MF implements automatic estimation of dictionary size and number of layers. Also, we provide a theoretical analysis of each operator and experimental validation of Deep MF, in Sections 3 and 4 respectively, by comparing with three other deep linear models, specifically, Deep SDL, Deep FICA and Deep NMF.

### 2.3 Deep Sparse Dictionary Learning

In the last decade, sparse dictionary learning (SDL), widely known as the algorithm Online Dictionary Learning (ODL) (Mairal et al., 2010; Liu et al., 2010), has been successfully applied to identify the concurrent BCNs of the human brain and the non-human primate brain from fMRI datasets (Lv et al., 2015; Zhang et al., 2018). In this category, to satisfy the requirements of hierarchical organization of BCNs, we propose a novel Deep SDL algorithm that is a multilayer extension of conventional shallow SDL-based methods. Briefly, fMRI signals from all voxels within the whole brain are extracted and are then organized as an extensive 2D matrix, where the number of columns represents the total brain voxels and the number of rows stands for the time points. For the first layer, the input 2D matrix is decomposed into the product of an incomplete/over-complete dictionary basis matrix (each atom representing a time series) and a feature matrix (representing this network’s spatial volumetric distribution). For each successive layer, the current features matrix is treated as an input matrix to be continuously decomposed. A particularly important characteristic of this Deep SDL framework is its ability to carry out over-complete decomposition for all layers; but, considering the finite features, the deep layers only concentrate on the incomplete decomposition.

If considering all layers, the optimization function of Deep SDL is:

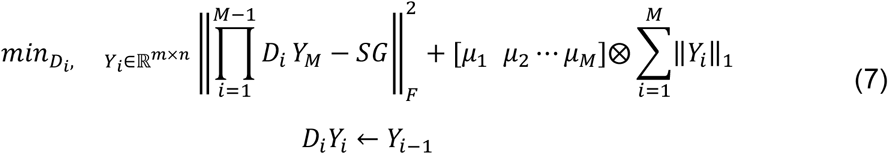

In Eq. (7), 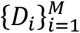 denotes the set of dictionaries; *i* represents the number of current layer, and *M* is the total number of layers, 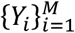 defines the set of hierarchical features, i.e., BCNs. And 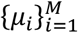 represents all sparsity trade-offs for all layers, respectively.

Deep SDL, like ODL, utilizes GD as the optimizer to update all parameters. Simple GD is an efficient optimizer, but only guarantees the convergence of convex problems. In Section 3.1, we also provide the requirements of GD to be a contraction operator. Briefly, the convergence property of GD heavily depends on the step length. The following equation shows the iterative format of simple GD:

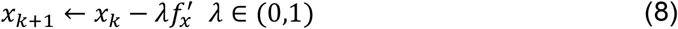

Compared with the ADMM optimizer used in Deep MF, the iterative function GD of Deep SDL is very simple. ADMM can utilize the current optima to update, and can be faster than GD, but ADMM requires that we can obtain the derivative of the optimization function.

### 2.4 Deep Fast Independent Component Analysis

ICA is a very popular and widely used data-driven computational technique, which was introduced to fMRI research over two decades ago (McKeown and Sejnowski 1998). In previous work, investigators have already reported that FICA using the Fixed-Point algorithm as an optimizer can be a very robust method (Hyvarinen, 1999). Inspired by Deep MF and FICA, the novel framework of Deep FICA aims to detect the hierarchically organized components. For simple shallow FICA, the original input matrix is decomposed as the weight matrix and independent component (IC) matrix. Applied to fMRI data, the ICs represent the BCNs. Similar to Deep MF and Deep SDL, in each layer of Deep FICA, the previous IC matrix is considered as the input signal matrix that will be decomposed using PCA and the Fixed-Point algorithm continuously (Figure 1). Deep FICA extracts only spatially independent features and can only solve the incomplete decomposition problem and not over-complete decomposition.

The optimization function of Deep FICA is:

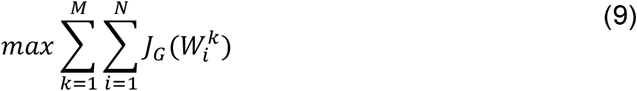

In Eq. (9), 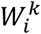 represents the *i*^th^ IC from the *k*^th^ layer. The maximum *J*_*G*_(∙) indicates the independency of each potential IC.

Considering each IC, Deep FICA and FICA both utilize an efficacy Fixed-Point algorithm to update the IC:

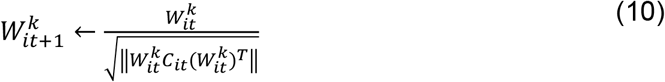

Compared with Deep MF and Deep SDL, Deep FICA is relatively easy to implement, since it does not include more complex algorithms, e.g., the RRO or the sparsity operator.

### 2.5 Deep Non-negative Matrix Factorization

Non-negative Matrix factorization (NMF) is a particularly useful family of techniques in data analysis. Before the wide utilization of the Deep CNN, NMF was a crucial technique to identify the features of a human face (Trigeorgis et al., 2016). In recent years, there has been a significant amount of research on deep factorization methods that focus on particular characteristics of both the data matrix and the hierarchical resulting factors. The application area of the family of NMF algorithms has grown significantly during recent years. It has been demonstrated that NMF can be a successful dimensionality reduction technique over a variety of application areas including, but not limited to, environmetrics, microarray data analysis, document clustering, face recognition and more. Moreover, due to its particular non-negative constraints, NMF can also be directly utilized to analyze the fMRI data/signal (Lee & Seung, 1999). Deep NMF provides an opportunity to detect the potentially hierarchical structures of BCNs.

Deep NMF focuses on the decomposition of the non-negative multivariate data matrix into hierarchical factors 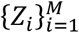 and 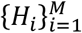, such that (Trigeorgis et al., 2016):

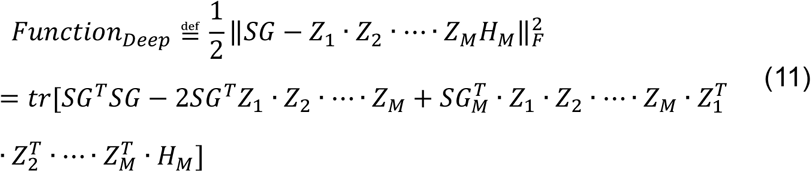

In Eq. (11), *SG* represents the input fMRI signal, and 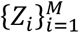 represents the weight matrix; 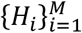 denotes the sets of non-negative components. *M* denotes the total number of layers. To calculate the optimal solutions of 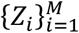 and 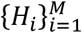 requires minimizing the loss function *Function*_*Deep*_ in Eq. (11). And *tr* represents the trace of the matrix.

A key difference between Deep NMF versus Deep SDL and Deep MF is the updating principle. Unlike Deep SDL and Deep MF, Deep NMF employs a fast policy to update these two factors: 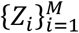 and 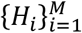 (Trigeorgis et al., 2016). This principle is shown as follows:

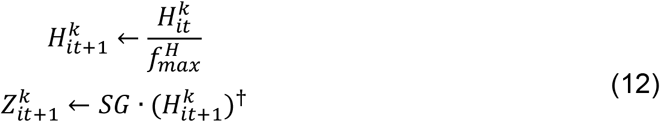

where 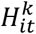 and 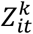 represent the non-negative components and weight matrix from the *k*^th^ layer, iteration number *it*. And operator (∙)^†^ represents the pseudo-inverse of the input matrix (Trigeorgis et al., 2016). The 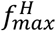 denotes the current maximum value of function *f*, related to 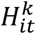.

Intuitively, these four deep linear models are each distinctive. Deep MF can be more intelligent, and automatically determine all hyperparameters. Deep SDL can perform over-complete decomposition for each layer. Deep FICA reveals the spatially independent and hierarchical components, and has a faster convergence velocity. Finally, Deep NMF could detect the non-negative components included in fMRI signals. We theoretically analyze the relative performance of each deep linear model in the following section.

## 3. Results: Theoretical Analyses

In this section, we employ mathematical theory, specifically real analysis, linear functional analysis and abstract algebra, to explain why different deep linear models have the distinctive characteristics that they do. In particular, we hope to explain:

i. The advantages of linear deep models over shallow models and deep nonlinear models;
ii. Why some deep linear models, e.g., Deep MF and Deep SDL, converge slowly while others, e.g., Deep NMF and Deep FICA, converge quickly;
iii. Why some deep linear models, e.g., Deep MF and Deep SDL, can better estimate connectivity strength, while others, e.g., Deep NMF and Deep FICA, can better estimate the spatial extent of connectivity networks.

### 3.1 Fundamental Interpretation of Each Linear Model

All theoretical analyses are based on the vital assumption that all mappings/operators/algorithms must be applied on a finite dimensional space. Please consult Appendix A for the mathematical details (Assumption 1.1, Lemma 1.1, Theorem 1.1). If considering any algorithm and/or process as an operator, Assumption 1.1 and Lemma 1.1 demonstrate that the norm of the operator should be equivalent, in order to dramatically simplify our discussion.

According to Theorem 1.1, if considering the shallow linear, deep linear and deep nonlinear models as approximations of the original function *f*(*x*), then, obviously, deeper models can employ more items such as 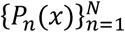 rather than just *P*(*x*). Thus, the deeper models can more accurately approximate the original function than a shallow model. Meanwhile, nonlinear models can have: 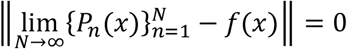. But to optimize the infinite items, it will be very time-consuming or even impossible to solve a non-polynomial (NP) complexity problem. Hence, the theorem also answers why nonlinear models require a sampling technique to reduce the complexity, e.g., Gibbs sampling for DBN (Hinton and Salakhutdinov, 2006; Hinton et al., 2012).

According to the discussion in the last section, we can abstractly describe each deep linear model using the combination of several operators. All operators involved in this study are given in Table 1.

**Table 1.**
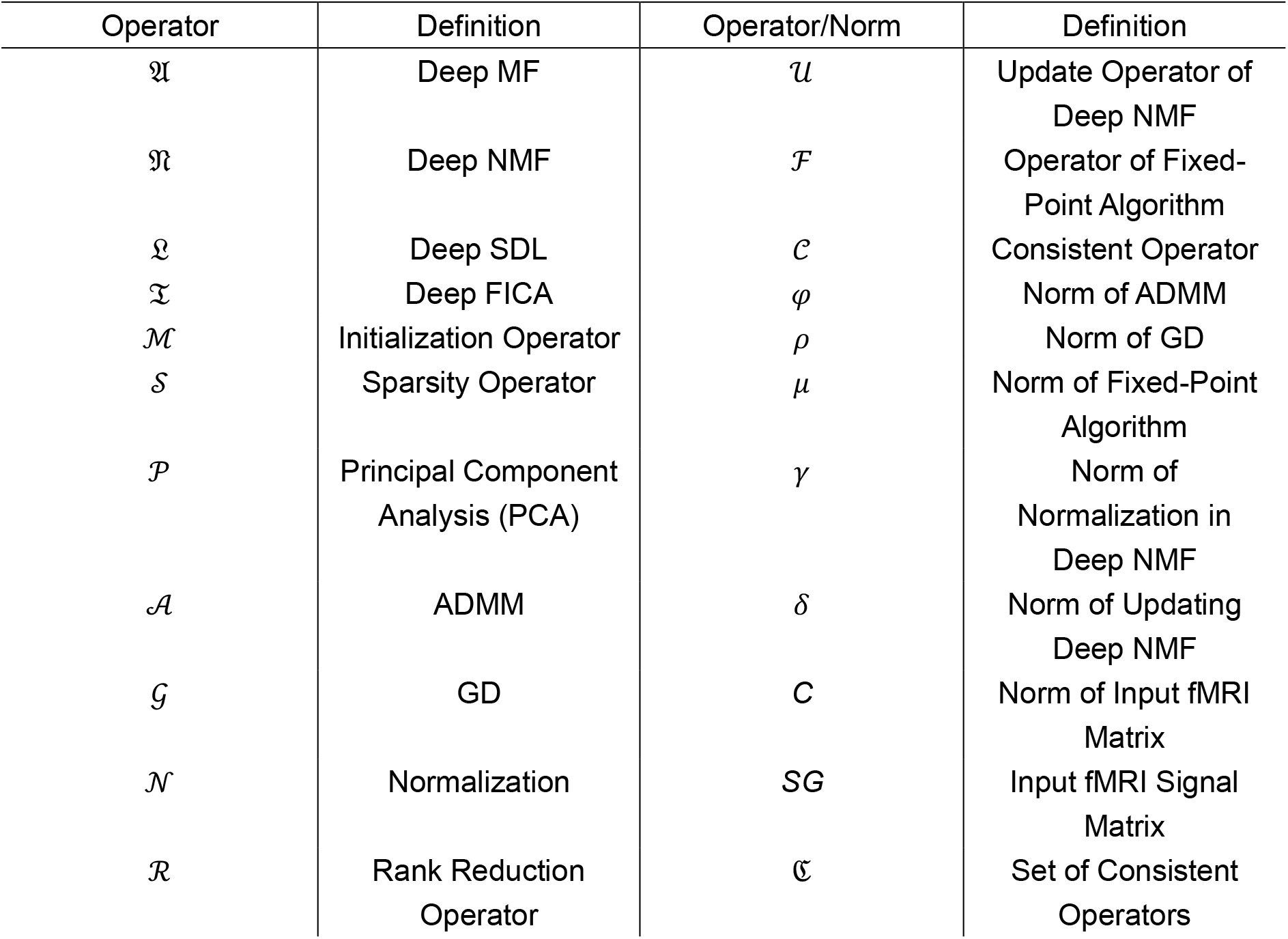
All definitions of operators and their norms.

In Table 2, we provide the definitions of sets involved in the following sections:

**Table 2.** All definitions of space and set

| Space/Set | Definition |
| --- | --- |
| $\mathbb{N}$ | Field of Natural Numbers |
| $\mathbb{R}$ | Field of Real Numbers |
| $\mathbb{K}$ | Field of Rational Numbers |

### 3.2 Intensity Similarity

As discussed in Sections 2.2 to 2.5, we have the following definitions:

***Definition 2.1*** If we denote Deep MF as an operator 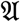, based on the description of Deep MF, considering the iteration *k*, we can denote 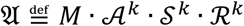.

***Definition 2.2*** If we denote Deep SDL as an operator 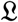, based on the description of Deep SDL, considering the iteration *k*, we can denote 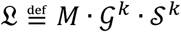.

***Definition 2.3*** If we denote Deep FICA as an operator 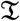, based on the description of Deep FICA, considering the iteration *k*, we can denote 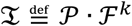.

***Definition 2.4*** If we denote Deep NMF as an operator 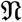, based on the description of Deep NMF, considering the iteration *k*, we can denote 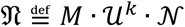.

According to Definitions 2.1 to 2.4, as well as Corollaries 1.2 to 1.3 and Theorems 2.1 to 2.10 as proved in Appendix B, using the inequality of norm, considering the iteration *k*, and let *SG* be the input matrix; for any operator applied on *SG*, we can derive the features as: *F*_1_, *F*_2_, *F*_3_, *F*_4_; then we have:

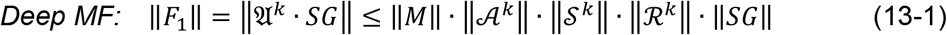

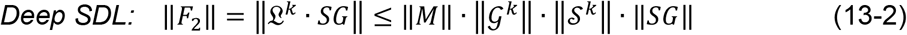

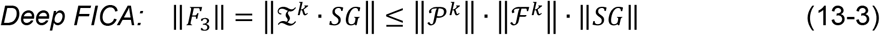

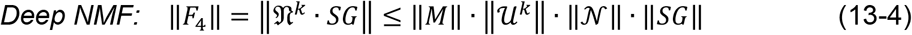

According to Lemma 1.1 and Theorems 2.1 to 2.6, operators 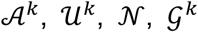, and 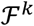 can be treated as contraction operators, which indicates that the norm of each operator should be larger than zero and smaller than one. Other operators are constant values, according to Theorems 2.1 to 2.10.

If we denote the norm of contraction operators as:

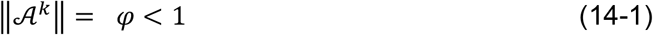

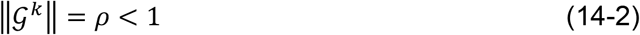

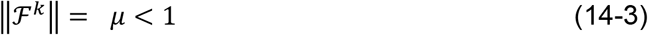

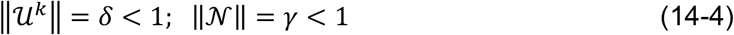

Despite the fact that the norms of operators 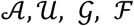 are not equivalent, according to Theorems 3.1 and 3.2 (see Appendix C), we consider an extreme condition *k* → ∞, and then we have: *φ* = δ = *ρ* = *μ* that indicates convergence to the global optimum. Then we can rewrite all equations 13-1 to 13-4 as:

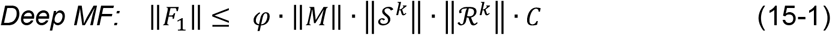

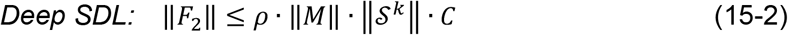

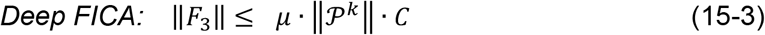

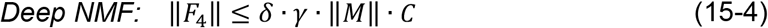

Obviously, based on Eqs. (15-1) to (15-4), we have the conclusion:

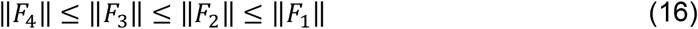

Since all features 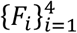 have the same dimensions, this inequality Eq. (16) can clearly explain why the intensity of features, i.e., the connectivity strength of voxels in the networks, varies based on the different models. In particular, *F*_4_ (the features derived from Deep NMF) should have the smallest intensity and *F*_1_ (the features obtained via Deep MF) should have the largest intensity. Meanwhile, Eqs. (15-1 to 15-4) also reveal the convergence velocity of each model. Since Deep MF contains the most operators with the complex optimization function ADMM, it should be slowest. Because Deep SDL uses a sparsity operator as well as GD, which is relatively slow, it is comparable in speed to Deep MF, even given a perfect step-length. Theoretically, Deep FICA and Deep NMF should have faster convergence.

### 3.3 Spatial Similarity

Spatial matching is another important way to measure the similarity between identified components and templates. To examine this property, we use Assumptions 3.1 to 3.2 and Lemma 3.1 to prove Theorem 3.1 in Appendix C:

#### Theorem 3.1

If we denote the following sets:

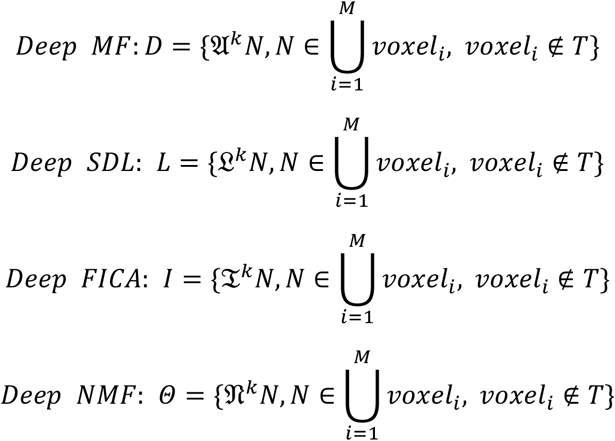

And considering the iteration *k*, and *k* > *K*, it implies:

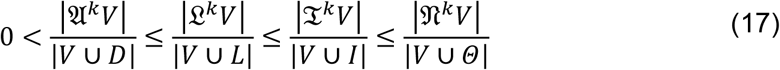

where |∙| denotes the number of positive elements. *Θ* represents the set that only contains the element 0. *T* represents the functional regions of brain.

Since the convergence of deep models is a vital issue when solving real world problems (Topol, 2019), Eq. (16) and Theorem 3.1 can explain the convergence of all deep linear models, considering enough iterations. Clearly, if we examine the spatial similarity between two BCNs, according to Theorem 3.1, we can conclude: with the same number of iterations, Deep NMF has the best performance on spatial matching, Deep FICA has the next best performance, and Deep SDL and Deep MF have the least. That is, the norm of the operator of Deep NMF is very small, and is iteratively applied on functional regions and background noise, which causes the intensity of functional areas to decrease very rapidly. However, since the intensity of background is very small, the noise can be reduced to near zero much faster than Deep MF and Deep SDL. The performance of Deep FICA on spatial matching should be comparable to Deep NMF; since a normalization operator is involved in Deep NMF, the intensity of components identified by Deep NMF should be smaller than Deep FICA. Theorem 3.2 included in Appendix C also explains that all deep linear models finally converge given enough iterations.

To test these theoretical analyses, in the next section, a simulated experimental reconstruction will be introduced as the ground truth templates for the first layer BCNs. These templates will be employed to construct the simulated fMRI signal and all deep linear models will be applied on the simulated data and their 1^st^ layer results will be compared to the templates. By examining the intensity similarity and spatial similarity to the ground truth templates, the correctness of the theoretical conclusions can be investigated.

## 4. Results: Experimental Validation

### 4.1 Simulated fMRI Data

In this work, we employ an *in silico* fMRI simulation method proposed previously (Zhang et al., 2018, 2019), using templates of BCNs (Smith et al., 2009) to test these proposed deep linear models. Specifically, we selected 12 BCNs (Table 3) that were originally derived using conventional shallow ICA (Smith et al., 2009).

**Table 3.** All abbreviations of BCNs in simulation

| Name/Number | Abbreviation | Name | Abbreviation |
| --- | --- | --- | --- |
| Primary Visual<br>Network/1 | VIS-1 | Auditory Network/7 | AUD |
| Perception Visual<br>Shape Network/2 | VIS-2 | Executive Control<br>Network/8 | ECN |
| Perception Visual<br>Motion Network/3 | VIS-3 | Left Frontoparietal<br>Network/9 | FP-L |
| Default Mode<br>Network/4 | DMN | Right Frontoparietal<br>Network/10 | FP-R |
| Brainstem &<br>Cerebellum Network/5 | B/C | Dorsal Attention<br>Network/11 | DAN |
| Sensorimotor<br>Network/6 | SM | Salience Network/12 | SN |

These template BCNs derived from resting-state fMRI have been released publicly and are considered to be functional brain areas covering a large part of cerebral cortex (Smith et al., 2009). Since all deep linear models should be evaluated equally using a known ground truth, we employ a simulation of resting-state fMRI signals. Following the fMRI simulation pipeline for spatially independent networks named Experiment 1 in the previous study (Zhang et al., 2019), the 12 templates (Smith et al., 2009) are collected as components/spatial features; and we adopt 12 time series, including 200 time points, derived from a previous study (Lv et al., 2015). The final simulation data is a matrix obtained as the product of time series and components.

The detailed parameters of each template are: 91×109 matrix, 91 slices, 2.0 mm isotropic voxels. The number of mask voxels is 262,309 and the number of time points is 200. All templates are registered to standard MNI space at 2.0 mm. This pipeline contains the steps of spatial artifact cleanup, distortions removal and cortical surfaces generation. After that, different subjects are aligned to the standard MNI space (Lv et al., 2015; Zhang et al., 2019).

Table 4 provides the main hyperparameter settings of the four proposed deep linear models, including: the number of components of the 1^st^ and 2^nd^ layers, the number of iterations and the step length of gradient descent, where applicable. Since the 1st layer may include noise components, we choose a larger number of components than the expected number of features, which in this case is at least a dozen ground truth template BCNs. For the 2^nd^ layer, which should have fewer high-level features, the number of components should be less than in the 1^st^ layer. Lv et al. (2015) introduce an experimental method to search for the best number of components, but, in fact, heuristically tuning the hyperparameters of deep models is very difficult. Since Deep MF is capable of estimating all these hyperparameters automatically, only the maximum number of iterations is given. For the other three methods, the hyperparameter values were chosen heuristically based on best matching to the ground truth templates for the 1^st^ layer and for perceived quality of the derived networks for higher layers.

**Table 4.** Important Hyperparameter Settings of Four Deep Linear Models

|  | Deep MF | Deep SDL | Deep FICA | Deep NMF |
| --- | --- | --- | --- | --- |
| Number of Components of 1<sup>st</sup> layer | N/A | 15 | 20 | 30 |
| Number of Components of 2<sup>nd</sup> layer | N/A | 13 | 10 | 15 |
| Number of Iterations | 100 | 100 | 100 | 100 |
| Step Length | N/A | 0.01 | N/A | N/A |

### 4.2 Investigating the First Layer Reconstructions of Each Deep Linear Model via Intensity, Spatial and Hausdorff Distances

We can quantitatively compare the identified components, i.e., BCNs, with the original ground truth, i.e., templates, in three distinct ways. First, the similarity can be calculated spatially, largely independent of the intensity of each voxel of the identified components. The definition of spatial similarity is:

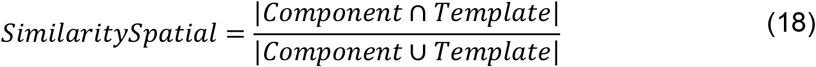

where |∙| represents binarization, which represent the voxels above a given intensity threshold. The spatial similarity is measuring the ratio of intersection and union of identified component and template.

In contradistinction, only considering the intensity of each voxel of the derived components, it is useful to calculate the distance between the intensities of components and templates. The definition of intensity similarity is:

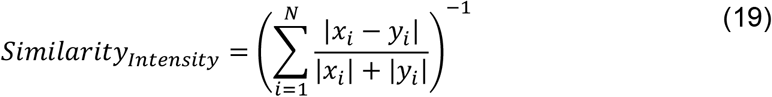

where |∙| represents the absolute value. Given a threshold, the intensity similarity is calculated via summed absolute value of intensity of component (denoted as *x_i_*) and template (denoted as *y_i_*) divided by the absolute value of their difference. *N* denotes the total number of voxels. Obviously, if all intensity values of identified component and template are equal, the intensity similarity approaches infinity.

Finally, to jointly consider both spatial and intensity matching, we use the Hausdorff Distance (HD):

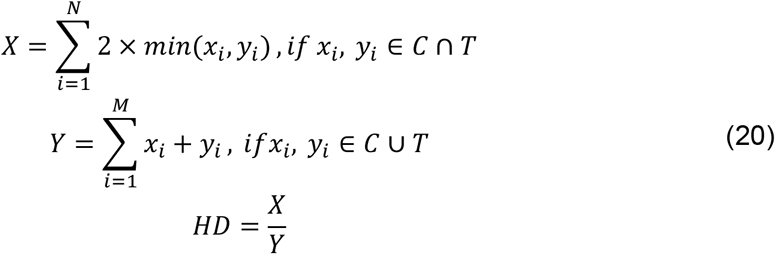

Briefly, *X* represents two times the minimum intensity value of intersection of component and template; and *Y* represents the summed intensity value of union of component and template. *C* and *T* represent the sets of components and templates, respectively. Therefore, HD includes the influences of intensity similarity and spatial overlap simultaneously.

The results show that Deep NMF and Deep FICA produce smaller network intensities than the templates, whereas Deep MF and Deep SDL yield larger intensities that better match the templates (Figure 2a). In contradistinction, there are generally more noisy areas detected from Deep MF and Deep SDL due to the larger norms of their iterative operators, compared to Deep NMF and Deep FICA (Figure 2b). Hence, Deep NMF and Deep FICA have better spatial similarity to the templates than the other two methods. This illustrates the trade-off between intensity matching and spatial matching. To view reconstructions for all 12 examined BCNs and for further details, please see Figure S1 included in the Supplemental Materials.

**Figure 2.**
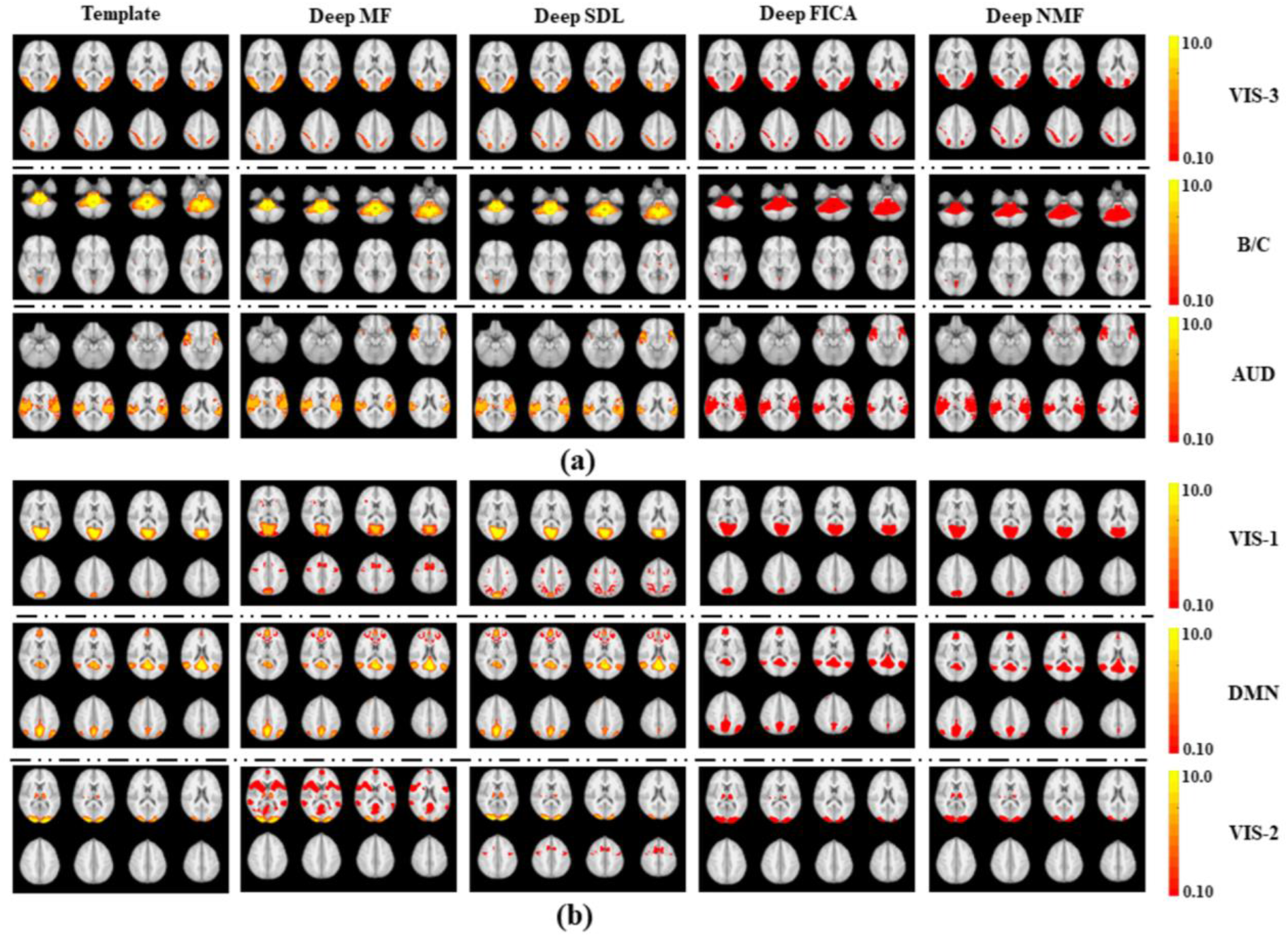
Comparison of six 1st layer networks from all four deep linear models with the ground truth templates from simulated fMRI data (see Table 3 for network abbreviations). The first column presents eight representative slices from each of six representative template networks. The second to fifth columns show the corresponding slices from the networks identified via Deep MF, Deep SDL, Deep FICA and Deep NMF, respectively. **(a)** The AUD, B/C and VIS-3 networks illustrate better intensity matching to the templates by Deep MF and Deep SDL than by Deep FICA or Deep NMF (see color bar of intensities measuring connectivity strength on the right). **(b)** The VIS-1, VIS-2 and DMN networks also show this same disparity among the deep linear models for intensity matching, but also show better spatial similarity to the templates for Deep FICA and Deep NMF compared to Deep MF or Deep SDL.

As defined by Eq. (18) to Eq. (20) in Section 4.2, the quantitative comparisons among the four deep linear models for intensity similarity, spatial similarity and the Hausdorff distance are provided by Figure 3. These quantitative results clearly demonstrate that Deep MF and Deep SDL provide the best intensity matching (Figure 3a), since their convergence velocity is relatively slow. Therefore, Deep MF and Deep SDL can reconstruct the most accurate connectivity strengths of each component from input fMRI signals, consistent with theory (Section 3). Considering spatial similarity, due to the fastest convergence velocity and non-negative normalization of Deep NMF, the intensity is reduced rapidly across iterations. Since the noise has smaller intensity than the signal, it is reduced much faster, which helps account for Deep NMF yielding the best spatial similarity results for most networks (Figure 3b). This result is also predicted by theory in Section 3. A rigorous proof is presented in the Appendices. It should be noted that Deep FICA has an inherent advantage over the other models for spatial matching since it is most similar to the shallow ICA analysis used to generate the ground truth templates for the BCNs.

**Figure 3.**
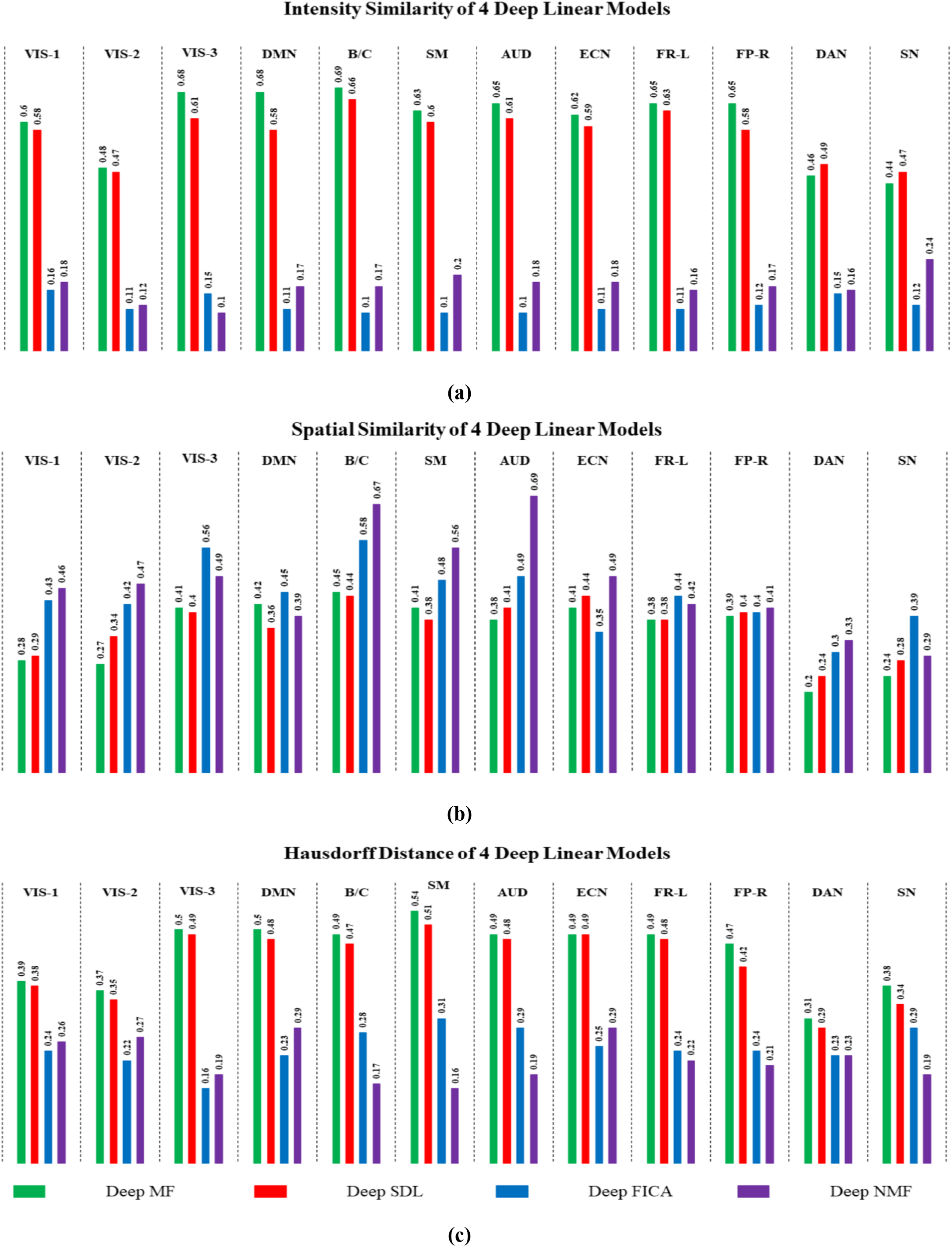
Comparisons of the twelve 1st layer networks of the four deep linear models for **(a)** intensity similarity to the ground truth templates; **(b)** spatial similarity to the ground truth templates; and **(c)** the Hausdorff Distance to the ground truth templates that jointly considers intensity and spatial similarity.

All proposed deep linear models can be evaluated by HD to consider both intensity and spatial similarity (Figure 3c). Deep MF generated the best performance for all 12 BCNs with Deep SDL running close behind. Hence, the additional RRO in Deep MF does yield advantages over the other three deep linear models. Similarly, the sparsity operator of Deep SDL and Deep MF help them outperform Deep FICA and Deep NMF, which both lack that capability.

### 4.3 The 2^nd^ Layer Networks of Each Deep Linear Model

Compared with the shallow 1^st^ layer features, it is difficult to successfully investigate the features of deeper layers because there is no widely accepted ground truth for those more complex higher-level networks. The 2^nd^ layer features can be comprehended as the recombination of 1^st^ layer features. Another challenge for testing deeper networks is that the deep linear models differ with regard to how many layers can be reconstructed from a given dataset. For example, Deep FICA can only decompose the simulated fMRI into two layers, but Deep MF can decompose the simulated signal into four layers. Given these constraints, as well as the altered connectivity strengths in the 2^nd^ layer relative to the 1^st^ layer, we limit the analysis of deeper networks to examining the spatial similarity between 2^nd^ layer networks of each of the four deep linear models with the shallow ground truth templates (Figure 4).

**Figure 4.**
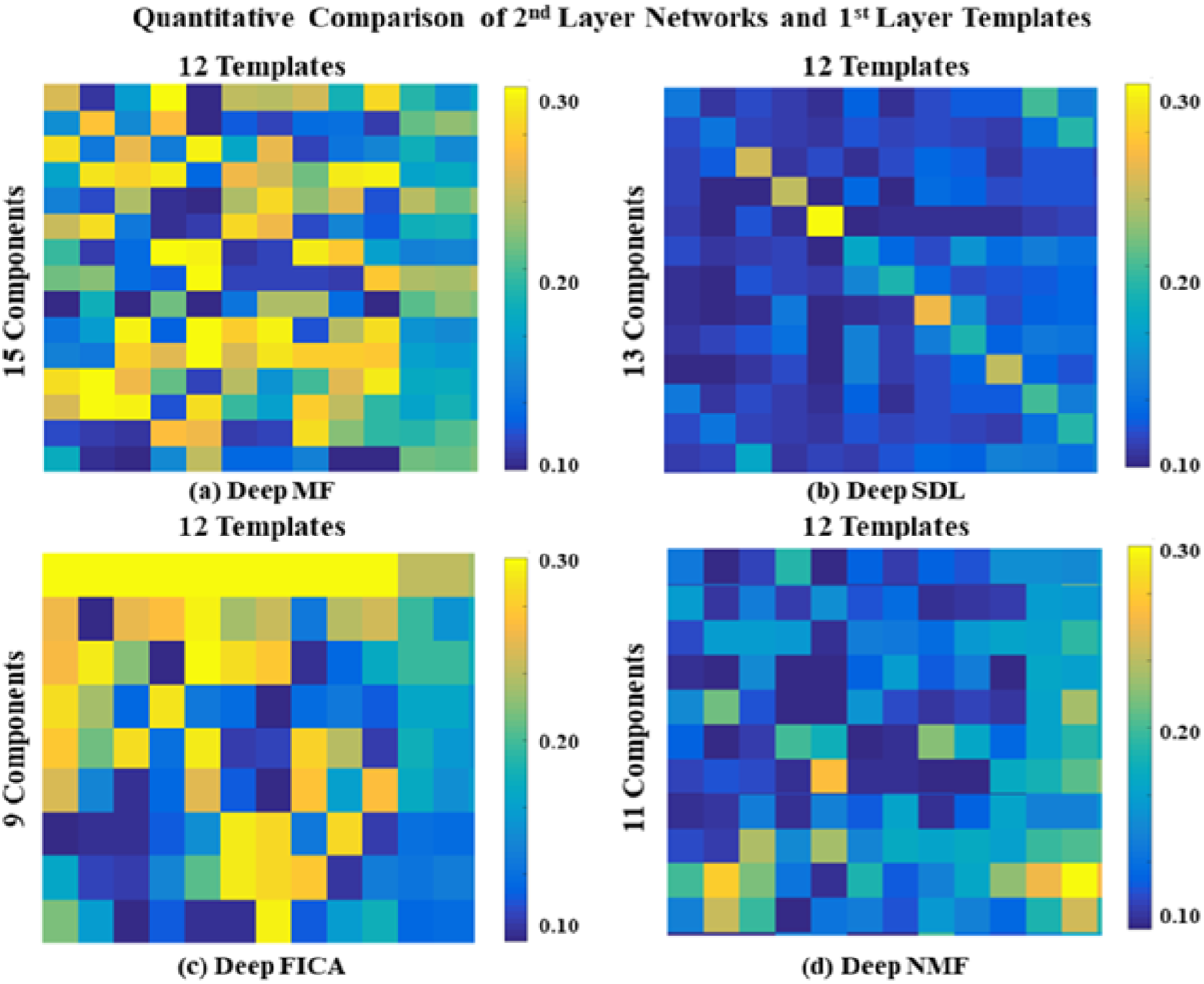
Comparisons of BCNs from 2^nd^ layer of Deep MF, Deep SDL, Deep FICA and Deep NMF. Each element represents the spatial similarity of the identified component and the ground truth templates; (a), (b), (c) and (d) are Deep MF, Deep SDL, Deep FICA and Deep NMF, respectively. The rows represent the identified 2^nd^ layer BCNs and the columns represent the ground truth templates of the simulated experiment.

Based on these preliminary comparisons, it is clear that the four deep linear models can produce different higher-level features. Most notable is that Deep SDL produces 2^nd^ layer networks that are the most spatially similar to the shallow ground truth templates, as shown by the larger main diagonal elements in its similarity matrix and the smaller off-diagonal elements (Figure 4b). Hence, Deep SDL does relatively little recombination of the 1^st^ layer features in its 2^nd^ layer. In contradistinction, the first component of the Deep FICA 2^nd^ layer (top row of its similarity matrix in Figure 4c) is very strongly correlated with 10 of the 12 ground truth templates and therefore appears to be a spatially “global” network. The 2^nd^ layer features of Deep NMF have overall the least spatial similarity with the ground truth templates (Figure 4d) whereas Deep MF produces the greatest variation in the correlations between its 2^nd^ layer features and the ground truth templates (Figure 4a).

Three representative 2^nd^ layer BCNs matched for each deep linear model are presented in Figure 5. The full set of non-noise 2^nd^ layer networks are given in Figure S2 of the Supplemental Materials. The top row of Figure 5 shows that the nodes of the ECN, including anterior cingulate cortex and medial prefrontal cortex, are represented in that 2^nd^ layer network for all four models. For both Deep MF and Deep FICA, the ECN is combined with nodes of the SN, including the insulae, pre-supplementary motor areas (pre-SMA), and premotor areas.

**Figure 5.**
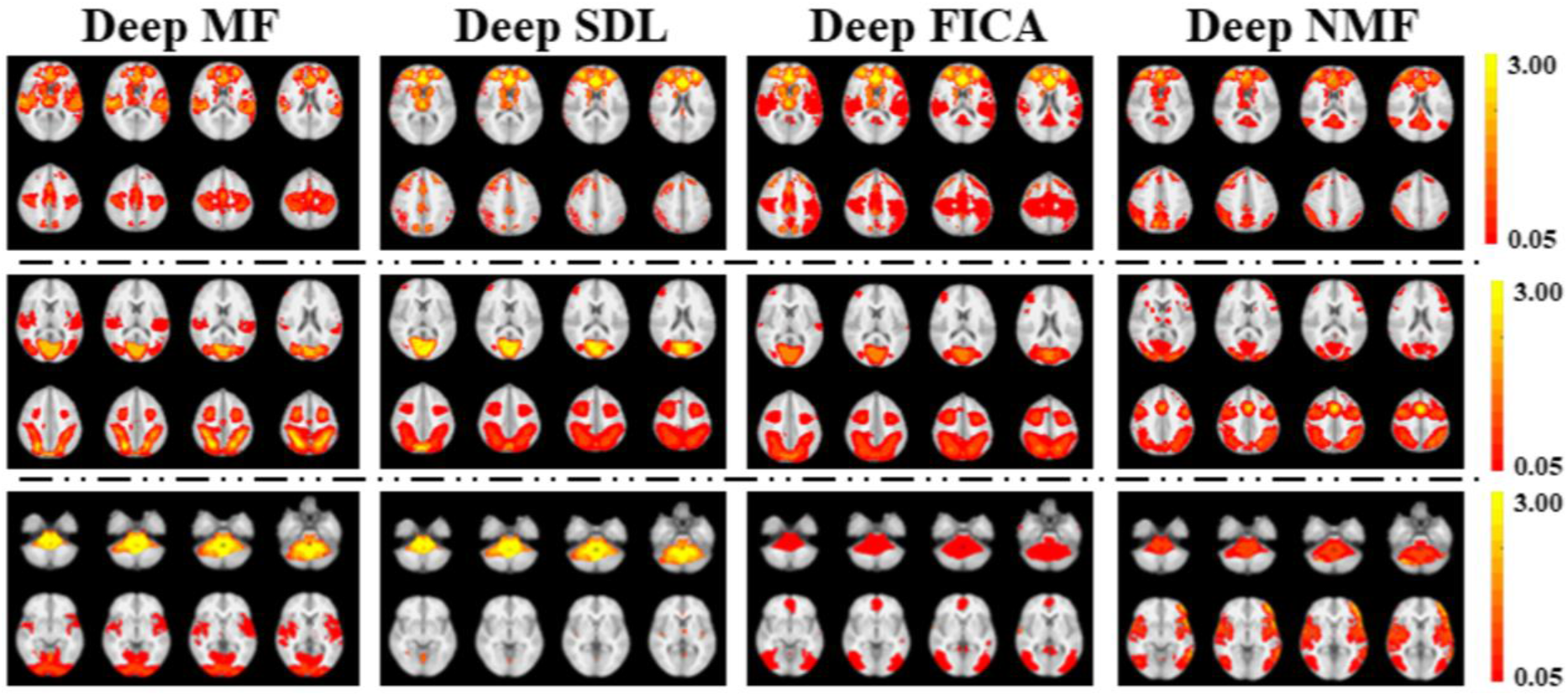
Comparisons of BCNs derived from the 2^nd^ layer of Deep MF, Deep SDL, Deep FICA and Deep NMF. Each column includes three representative 2^nd^ layer networks from a deep linear model, matched across models in each row.

For Deep NMF and Deep SDL, however, the ECN is joined instead with nodes of the DMN, including the precuneus, posterior cingulate cortex and the superior parietal lobules. This higher-level connectivity between ECN and DMN is weaker for Deep SDL than Deep NMF, whereas connectivity within ECN is stronger for Deep SDL than Deep NMF, in keeping with the observations from Figure 4 that Deep SDL preserves 1^st^ layer networks the most of all four algorithms, whereas Deep NMF preserves 1^st^ layer networks the least (Figures 4 & S2). It can be observed that the same 2^nd^ layer network of Deep FICA also contains parts of the DMN, most notably the posterior cingulate cortex, although to a lesser extent than Deep NMF. Therefore, Deep FICA recombines nodes of three different 1^st^ layer spatially independent components (DMN, ECN & SN) into a single 2^nd^ layer independent component. Figure S3 of the Supplementary Materials provides a spatial similarity matrix for the non-noise 2^nd^ layer networks for Deep SDL, Deep FICA and Deep NMF with reference to Deep MF.

The middle row of Figure 5 shows that all four models produced a 2^nd^ layer network consisting of VIS-1 and DAN. However, Deep NMF additionally included parts of VIS-2, VIS-3 and pre-SMA whereas Deep MF additionally included VIS-3 and posterior perisylvian regions. The bottom row of Figure 5 illustrates links of the brainstem and cerebellum with visual cortex. However, Deep MF finds correlations of B/C with VIS-1 & VIS-2 whereas Deep FICA and Deep NMF finds correlations with VIS-3 instead. Both Deep MF and Deep NMF include perisylvian regions in this 2^nd^ layer network as well. Given the relatively slow convergence of Deep SDL via gradient descent, these features might be expected in its 3^rd^ layer instead.

We also compare computation time for the four deep linear models presented in this work on our computing cluster (Figure 6), which demonstrates that Deep FICA is the fastest and Deep SDL is by far the slowest. Deep MF provides the best trade-off between speed and performance as judged by reconstruction accuracy for 1^st^ layer networks (Figure 3).

**Figure 6.**
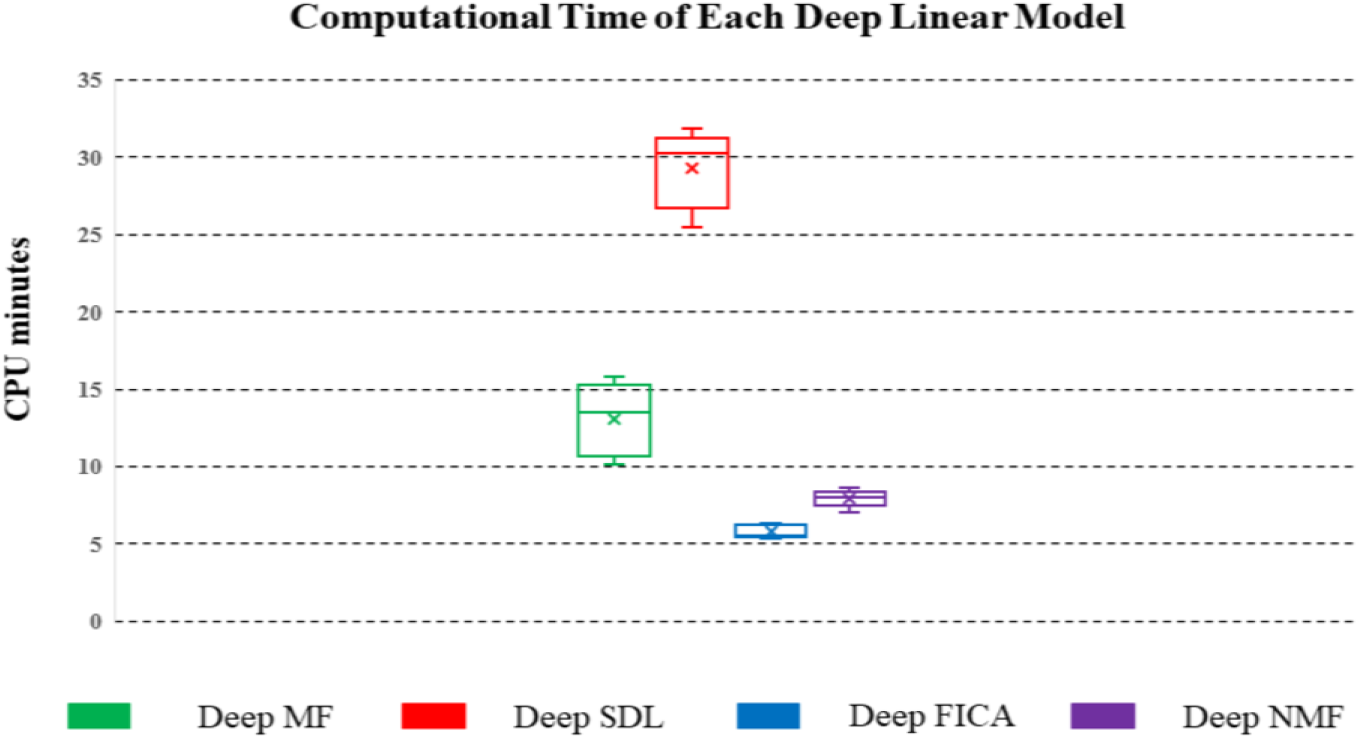
Comparisons of computation time of 10 independent runs, using the same number of iterations and the same simulated fMRI dataset for Deep FICA (blue), Deep SDL (red), Deep MF (green) and Deep NMF (purple). The box plots give the mean and standard deviation of the CPU time in minutes for the 10 runs.

## 5. Discussion

We have introduced novel deep linear models that integrate multiple operators to extract hierarchical spatial features in fMRI data. These models bridge the gap between traditional shallow linear models (Andersen et al., 1999; Beckmann et al., 2005; Calhoun et al., 2001; Hyvarinen, 1999; Lee et al., 2011; Lee et al., 2016; Mairal et al., 2010; McKeown & Sejnowski, 1999) and newer deep nonlinear models (Hu et al., 2018; Huang et al., 2018; Dong et al., 2020; Zhang et al., 2020). The primary advantages of the proposed algorithms over more complex deep nonlinear models are to quickly and easily map the hierarchical organization of BCNs without requiring large amounts of fMRI data or HPC clusters with GPUs or TPUs. The behavior of deep linear models is also more explainable than are, for example, CNNs and DBNs, as we show through theoretical predictions of their relative performance (Section 3) that are validated via simulations (Section 4). Furthermore, convergence to the global optimum can be guaranteed for deep linear models with convex optimization functions, unlike deep nonlinear models where such convergence is rarely achieved in practice. This is important given the recent realization that real-world imaging applications often suffer from underspecification, resulting in wildly unpredictable performance from any particular deep nonlinear network due to convergence to different local optima from different random initial conditions despite identical training data and hyperparameters (D’Amour et al., 2020).

Deep MF employs ADMM, which is a distributed optimization algorithm particularly well suited to compositional analysis of hierarchical modular systems, and also utilizes RRO for data-driven determination of all hyperparameters, which can be considered an intelligent factorization method. This is a major advantage over many conventional shallow data-driven fMRI connectivity reconstruction methods and the other three deep linear models presented here, as well as more complex deep nonlinear models, all of which must be manually tuned for hyperparameter settings. Deep SDL can explore more potential components than other models, even more than the number of original time points, via over-complete decomposition. Deep NMF converges very rapidly and recognizes the non-negative constraints of BCNs in fMRI. Finally, Deep FICA efficiently maps hierarchically spatially independent BCNs and is even easier to implement than the other peer deep linear models, especially given the wide usage of shallow ICA models for unsupervised fMRI mapping.

In this research, we also introduce an innovative framework for studying the relative performance of deep linear models, both theoretically by comparing their mathematical structure as well as *in silico* via fMRI simulations. Evaluating the 1^st^ layer reconstructions of these deep linear models using simulated fMRI data from widely accepted ground truth BCNs, we find that Deep MF and Deep SDL are clearly superior for computing connectivity strength whereas Deep NMF and Deep FICA are modestly better for mapping spatial extent. These results were predicted from the unique mix of mathematical operators used in each of the four methods (Section 3). Overall, Deep MF provided the most robust combination of intensity matching, spatial matching and computational efficiency of all four techniques. This can be attributed to its joint use of sparsity and rank reduction operators in conjunction with the distributed ADMM optimization function. We also discovered that deeper features such as the 2^nd^ layer BCNs are recombinations of the 1^st^ layer networks and that these can vary among the four deep linear models. For example, Deep SDL produces the least recombination of the 1^st^ layer networks in its 2^nd^ layer. This can be attributed to its relatively slow convergence velocity using gradient descent optimization; therefore, more low-level network recombination is seen in its 3^rd^ layer instead. Another important factor that differs among the deep linear models is the number of spatial features that can be accommodated at each level of the hierarchy and the maximum number of layers for any given dataset. For example, Deep FICA supports the fewest number of meaningful components at the 2^nd^ layer (nine) and only two layers total; therefore, its 2^nd^ layer networks would be the least sparse. This can be seen in the top row of Figure 5 in which Deep FICA combines nodes of DMN, ECN and SN into a single 2^nd^ layer network, unlike the other models that only incorporate two of the three 1^st^ layer networks, but which can instead generate even deeper networks beyond the 2^nd^ layer. Hence, the choice of deep linear model matters for exploring higher-level BCNs. The mathematical evaluation framework and the fMRI simulation procedure provided in this work should enable further development of deep linear models that are optimized for different types of real-world applications in biomedical imaging, with Deep MF as the current best algorithm for fMRI hierarchical functional connectivity mapping.

One shortcoming of the current work is that the ground truth templates for testing the 1^st^ layer networks were generated using conventional shallow ICA (Smith et al., 2009), which is currently the most widely accepted technique for data-driven analysis of functional connectivity. Aside from giving Deep FICA an inherent advantage, these spatially independent BCNs do not adequately evaluate the ability to reconstruct overlapping networks that is a property of methods such as shallow or Deep SDL. We also do not comprehensively investigate the properties of the deeper layers of these four models, which is an extensive topic that is beyond the scope of this paper, especially considering the absence of gold standards for these more complex high-level networks as well as the wide variation among deep linear models in key attributes such as convergence velocity and enforcement of sparsity.

In this initial exploratory work, many of the derived 2^nd^ layer BCNs demonstrate neurobiological face validity. For example, the SN is known to modulate the anticorrelated connectivity of the DMN and the ECN (Menon & Toga, 2015); hence the linkage of their nodes into a single higher-level network (Figure 5, top row). The functional coupling of vision networks with the DAN shown in Figure 5 (middle row) is also well known, given the role that the DAN plays in visual attention and eye movements (Vossel et al., 2014). Future neuroscientific studies will be required to empirically validate the deep features of these models using demographic, clinical, cognitive, behavioral and/or electrophysiological data.

Since these deep linear models do not require large training datasets nor specialized computing infrastructure, they can be easily applied to clinical research with the potential to generate novel functional connectivity biomarkers of neurodevelopmental, neurodegenerative, and psychiatric disorders (Parkes et al., 2020), including for diagnosis, prognosis and treatment monitoring. This is particularly significant given the recent observation that neuropathology and psychopathology often affect low-level network connectivity differently than high-level network connectivity. For example, many different psychiatric disorders have been found to decrease lower-order sensory and somatomotor network connectivity in a uniform manner across patients (Elliott et al., 2018; Kebets et al., 2019), while increasing distinctiveness among patients in networks at higher levels of the hierarchy (Kauffman et al., 2017; Parkes et al., 2020). In fMRI studies of mild traumatic brain injury (TBI), altered functional connectivity has been found early after concussion both within individual BCNs, such as the SN, DMN and ECN, as well as between different BCNs (Palacios et al., 2017). Interactions of BCNs, such as that of the SN with the DMN, are thought to be especially important for outcome after TBI and can be used to guide personalized treatment (Jilka et al., 2014; Li et al., 2019). Disordered coupling of the SN with the DMN and ECN has also been shown in mild cognitive impairment (Chand et al., 2017). Hence, prevalent neurological disorders such as head trauma and neurodegenerative disease are thought to affect multiple levels of the human brain’s hierarchical organization. Such high-level interactions between DMN, ECN and SN can be investigated with deeper layers of these hierarchical linear models that integrate their spatially distinct gray matter nodes into a single larger-scale network, as seen in Figure 5 (top row). These examples show how more principled data-driven characterization of this hierarchy, particularly at its higher levels, holds great promise for providing clinically actionable biomarkers of neurological and psychiatric diseases.

The benefits of deep linear models gain in importance as the spatial and temporal resolution and sensitivity of fMRI continue to increase with improved MR imaging hardware and pulse sequences, e.g., the advent of SLice Dithered Enhanced Resolution Simultaneous MultiSlice (SLIDER-SMS) imaging (Vu et al., 2018) and MultiBand MultiEcho (MBME) imaging (Boyacioğlu et al., 2015; Cohen et al., 2020). Higher fMRI sensitivity and spatial resolution will enable mesoscale functional imaging that supports more 1^st^ layer components of deep linear models to uncover subnetworks of the BCN templates used in this work. This will also permit the use of deeper models to extract more levels of the hierarchy of functional connectivity. Whereas many widely used methods for performing time-varying fMRI analysis are heuristic rather than data-driven, such as those with arbitrary time windows (Iraji et al., 2020), advances in fMRI temporal resolution can be combined with deep linear models that perform joint spatiotemporal decomposition for principled unsupervised dynamic functional connectivity mapping that reveals ever more of the human brain’s hierarchical organization.

## Abbreviations

ADMM: Alternating Direction Method of Multipliers
BCN: Brain Connectivity Network
CNN: Convolutional Neural Network
CPU: Central Processing Unit
DBN: Deep Belief Network
DCAE: Deep Convolutional Auto Encoder
Deep FICA: Deep Fast Independent Component Analysis Deep
MF: Deep Matrix Fitting
Deep NMF: Deep Non-negative Matrix Factorization Deep
SDL: Deep Sparse Dictionary Learning
DNN: Deep Neural Network
fMRI: Functional Magnetic Resonance Imaging
GD: Gradient Descent
GLM: General Linear Model
GPU: Graphics Processing Unit
HD: Hausdorff Distance
ICA: Independent Component Analysis
IS: Intensity Similarity
LASSO: Least Absolute Shrinkage and Selection Operator
RBM: Restricted Boltzmann Machine
RRO: Rank Reduction Operator
rsfMRI: Resting-State Functional MRI
SS: Spatial Similarity
tfMRI: Task-Evoked Functional MRI
TBI: Traumatic Brain Injury
TPU: Tensor Processing Unit

## 6. Acknowledgements

This work was supported by the U.S. National Institutes of Health [R01 MH116950, U01 EB025162] and U.S. Department of Defense [W81XWH-14-2-0176].

## Appendix A

***Assumption 1.1*** For any operator discussed in this study, we have: 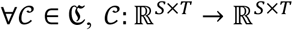. This assumption demonstrates that all operators are mapping from the finite dimensional space to another finite dimensional space, which is also reasonable in the real world.

### Lemma 1.1 (Norm Equality)

Given any arbitrary norm ‖∙‖ and/or their finite linear combination 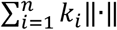 denoted based on any finite set, this norm or their finite linear combination is equivalent to ℓ_2_ norm (e.g., ‖∙‖_2_).

***Proof***: We denote ℓ_1_ and ℓ_2_ norm in finite dimensions, such as:

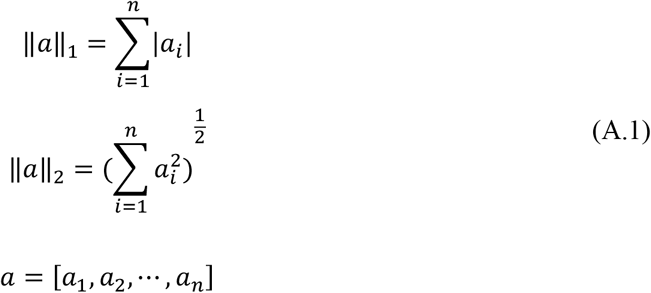

Obviously, since all norms are non-negative, according to Eq. (A.1), we have:

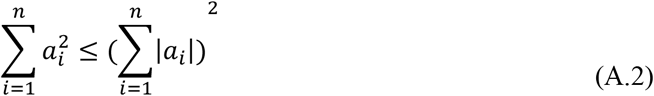

Eq. (A.2) implies:

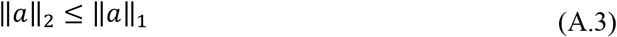

And, based on Cauchy-Schwarz inequality, we have:

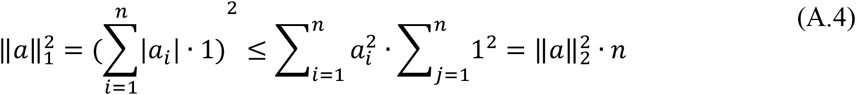

It implies:

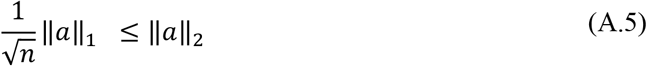

According to the theorem of norm equality (Rudin, 1973), given an arbitrary finite dimensional space, if and only if the following inequality holds:

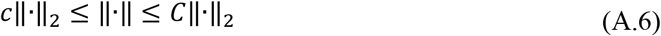

Thus, the norm ‖∙‖ is equivalent to ‖∙‖_2_. Since Eq. (B.3) and Eq. (B.4) hold, we have:

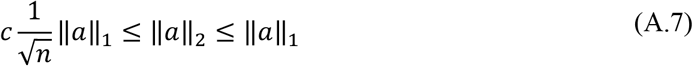

It implies ‖∙‖_1_ and ‖∙‖_2_ are equivalent. Similarly, we can prove 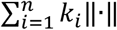 is also equivalent to ‖∙‖_2_.

### Theorem 1.1 (Superiority of Deep Linear Models)

Given a real function *f*(*x*) and *m*({*x* ∈ [a,b]:|*f*(*x*)|=±∞})=0. If considering the series of polynomials 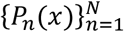, we have: if *N* is large enough, we have; if *N* → ∞ 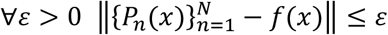, we have: 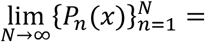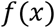 however, for any shallow model, since *N* should be bounded, we only have: 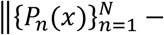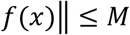.

***Proof***: According to 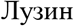 (Luzin) Theorem (Royden, 1968), we have a close set:

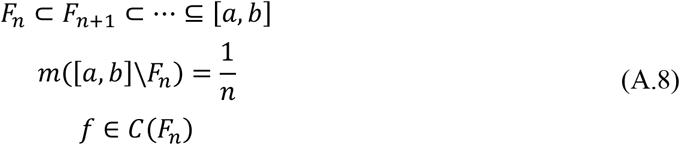

Then we have a consistent real function *g*(*x*), and obviously we have:

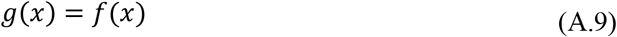

Since for any continuous real function, we have:

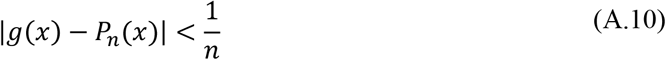

Let 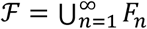, and obviously we have:

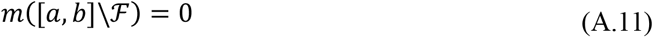

If 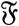 is a real function denoted on set 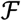, it indicates:

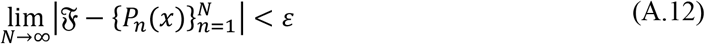

then we have 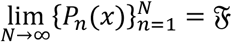,

Moreover, it is easy to prove 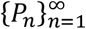 denoted on [*a*, *b*] as a ring 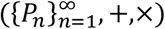 (Dummit, 2004; Kadison, 1997). And *m*(∙) represents a Lebesgue measure.

### Lemma 1.2 (Contraction of Operators Combination)

Given two contraction mappings Φ_1_ and Φ_2_, we have the composite of two contraction mapping as Φ_2_ ∙ Φ_1_. The composite mapping Φ_2_ ∙ Φ_1_ must be contractive.

***Proof***: According to the definition of contraction linear operator, we have:

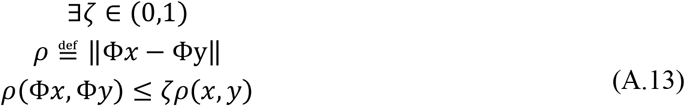

Obviously, and we have:

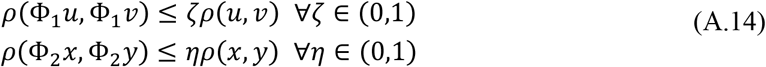

If we set:

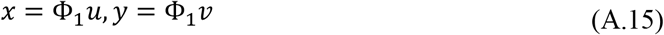

the inequality below holds:

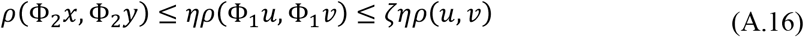

Since the definition as

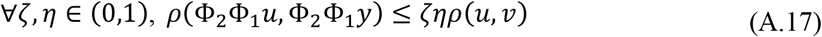

### Corollary 1.1 (General Contraction Operator)

According to Lemma 1.2, if denote the operators 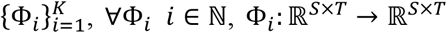; considering any combination of operators: Φ_*K*_ ∙ ⋯ ∙ Φ_2_ ∙ Φ_1_, if at least a single operator Φ_*i*_ is contraction operator, and other operators are bounded, such as ∀*i* ≠ *k* ‖Φ_*i*_‖ ≤ *M*. If and only if 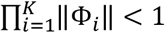, the combination of operator series Φ_*K*_ ∙ ⋯ ∙ Φ_2_ ∙ Φ_1_ is a contraction operator.

***Proof***: Obviously, according to Lemma 1.2, use a series as 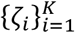 to replace ζ, η ∈ (0,1),

Obviously, we have:

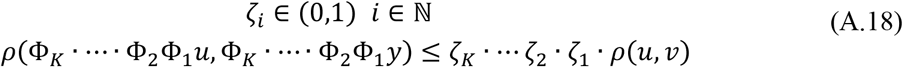

Since ζ_*K*_ ∙ ⋯ ζ_2_ ∙ ζ_1_ < 1, we have proved this corollary.

### Corollary 1.2 (Iterative Contraction Operator)

According to Lemma 1.2, if denote the operators 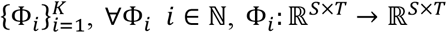; considering any combination of operators: Φ_*K*_ ∙ ⋯ ∙ Φ_2_ ∙ Φ_1_, if at least a single operator Φ_*i*_ is contraction operator, and other operators are bounded, such as ∀_*i*_ ≠ *k*, ‖Φ_*i*_‖ ≤ *M*. If and only if 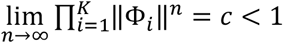, the combination of operator series 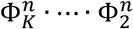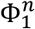

***Proof***: Obviously, according to Lemma 1.2 and Corollary 1.1 and 1.2, use a series as 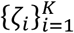 to replace ζ, η ∈ (0,1),

And we have:

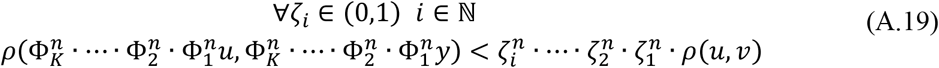

Since 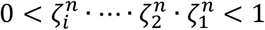, we have proved this corollary.

## Appendix B

***Definition 2.1*** If we denote Deep MF as an operator 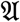, based on the description of Deep MF, considering the iteration k, we can denote 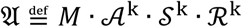.

***Definition 2.2*** If we denote Deep SDL as an operator 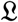, based on the description of Deep SDL, considering the iteration k, we can denote 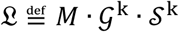.

***Definition 2.3*** If we denote Deep FICA as an operator 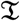, based on the description of Deep FICA, considering the iteration k, we can denote 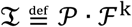.

***Definition 2.4*** If we denote Deep NMF as an operator 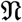, based on the description of Deep NMF, considering the iteration k, we can denote 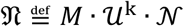.

### Theorem 2.1 (Contraction of ADMM Operator)

ADMM could be considered as contraction operator. It can be treated as a general iterative contraction operator in finite dimensionality space. We have 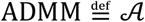. If denote the 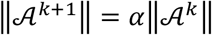, and *β* should be step length, i.e., penalty parameter, if *n* → ∞ 0 < (*αβ*)^*n*^‖*BN*‖ < 1, 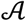 can be considered as a contraction operator. And ‖*BN*‖ denotes the norm of different residual error, considering two distinctive input matrices.

***Proof**: X* and *Y*, represent the two input matrices.

Consider the iterative format of ADMM as

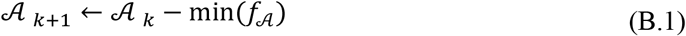

And it also can imply:

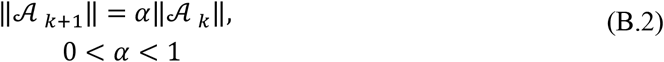

According to the definition of contraction operator, we have:

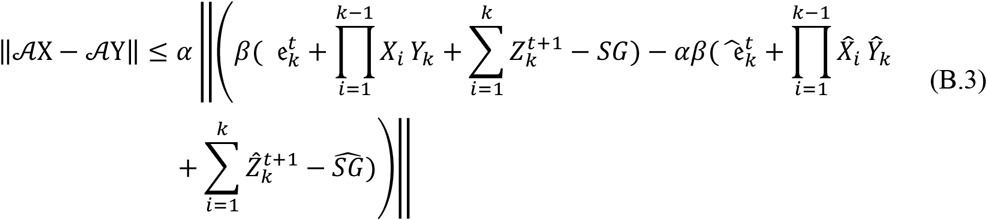

And we also have:

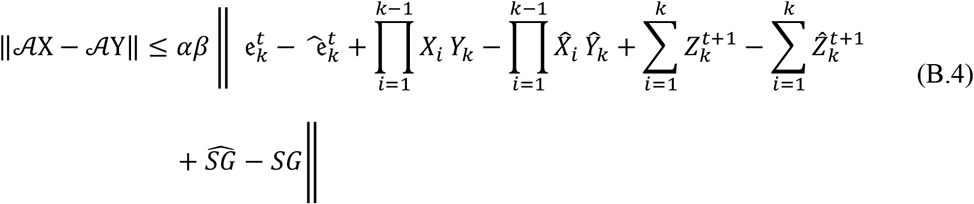

Since 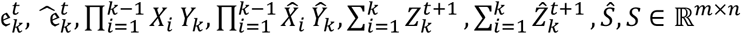, they are obviously bounded; and using Corollary 1.1 and 1.2, we have:

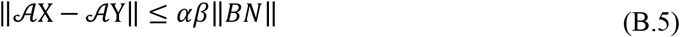

Obviously, it demonstrates:

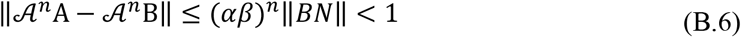

If and only if 0 < (*αβ*)^*n*^ < 1, or 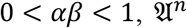 is equivalent to a contraction operator. According to Lemma 1.2 and Corollary 1.1, 1.2, it also indicates: when *n* is large enough, *n* > *N*, we have:

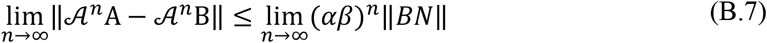

Obviously, if and only if 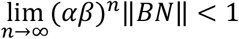, the iterative ADMM operator can be equivalent to a contraction operator.

### Theorem 2.2 (Initialization Operator is bounded)

If we denote the sparse operator as 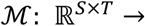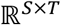, we have 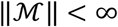.

***Proof***: according to the definition of operator norm (Rudin 1973), 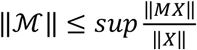; obviously, 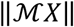 and ‖*X*‖ is bounded, since both of norms are based on finite dimensional matrix. And if we denote:

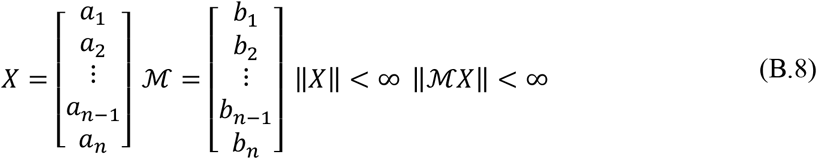

Obviously, 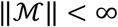.

### Theorem 2.3 (Sparsity Operator is bounded)

If we denote the sparse operator as 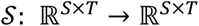, we have 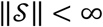.

***Proof***: according to the definition of operator norm (Rudin, 1973), 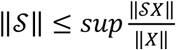; obviously, 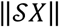 and ‖*X*‖ is bounded, since both of norms are based on finite dimensional matrix. And if we denote:

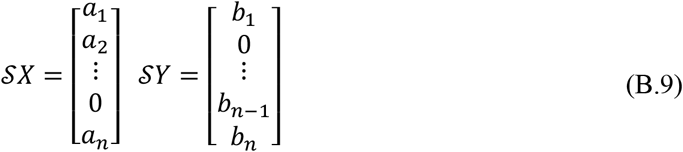

and we examine:

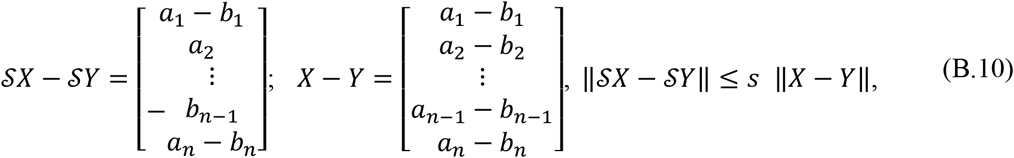

Without loss of generality, and based on Lemma 1.2, we calculate the ℓ_2_ norm, and we have:

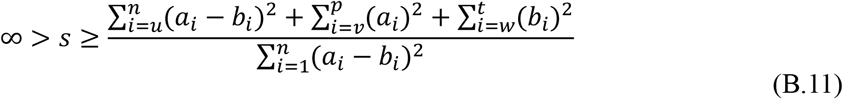

This inequality demonstrates that 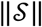 is a bounded. And 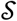 is a bounded operator.

### Theorem 2.4 (Rank Reduction Operator is bounded)

If we denote the sparse operator as 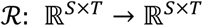, we have 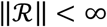.

***Proof***: According to the definition of operator norm (Rudin, 1973), 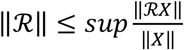 obviously, 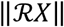 and ‖*X*‖ is bounded, since both of norms are based on finite dimensional matrix. And if we denote:

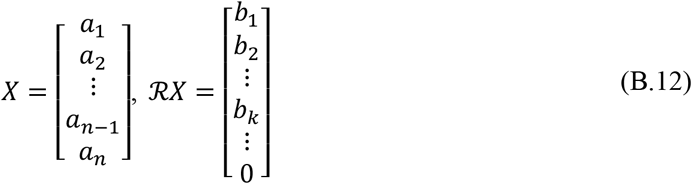

Eq. (B.12) implies:

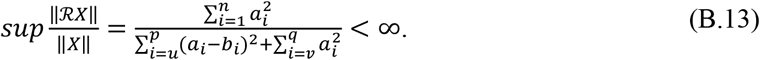

### Theorem 2.5 (Normalization Operator of Deep NMF is bounded)

If we denote the normalization operator of Deep NMF as 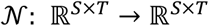, we have 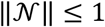.

***Proof***: according to the definition of operator norm (Rudin, 1973), 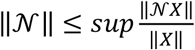; obviously, 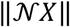 and ‖*X*‖ is bounded, since both of norms are based on finite dimensional matrix. And if we denote:

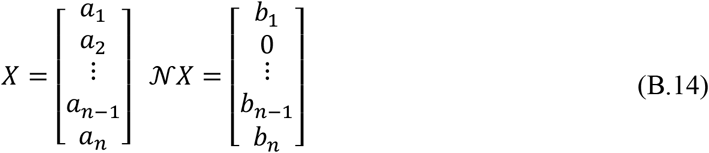

According to Eq. (B.14), we need to notice: 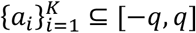, 1 ≤ *q* < ∞; 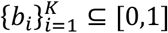.

Obviously, 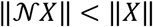. Finally, we have: 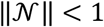.

### Theorem 2.6 (Contraction of Updating Operator Deep NMF)

If we denote the updating operator as 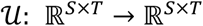, we have 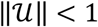.

***Proof***: according to the definition of operator norm (Rudin, 1973), 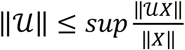; obviously, 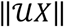 and ‖*X*‖ is bounded, since both of norms are based on finite dimensional matrix. And if we denote:

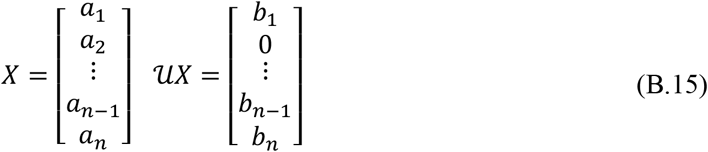

According to the iterative format of Deep NMF, we need to notice: 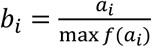; obviously, 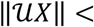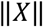. Finally, we have: 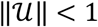. Otherwise, if 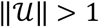, when *k* → ∞, we have: 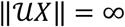.

### Theorem 2.7 (Contraction of GD Operator)

Gradient Descent (GD) is a bounded contraction operator, if and only if the derivative of target function is bounded: 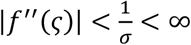is the step length.

***Proof***: The standard iteration format is:

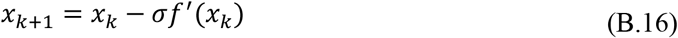

Using the definition of operator, we have:

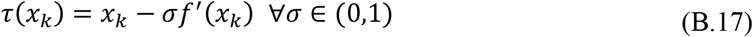

And we have:

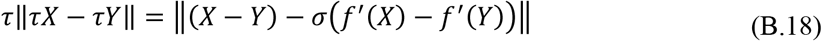

Using Mean value theorem, we have:

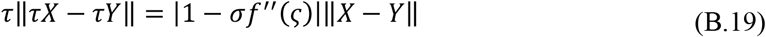

According to the definition of contraction operator (Rudin, 1973), if and only if:

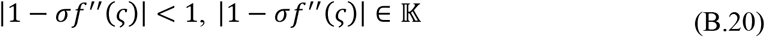

It also implies, w1hen the following inequality holds:

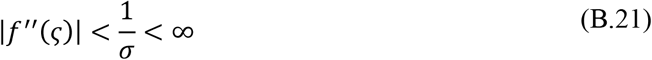

GD is considered as a contraction mapping/operator. Without generality, we can set 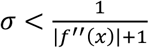.

And obviously, using multiplicative inequality, we have:

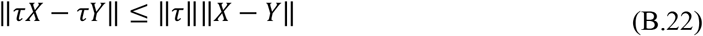

Since *X* and *Y* both denote in finite ℓ^2^ space, we have:

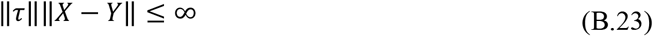

Using Uniformly bounded theorem, we have:

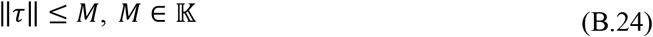

GD is a bounded mapping/operator.

According to Lemma 1.2, and Corollary 1.1-1.2, obviously, for *n* iterations for an operator, and if we set the accuracy level as *ɛ*, we have:

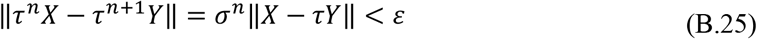

Since *X* and*Y* is both denoted in finit ℓ^2^ space, we have:

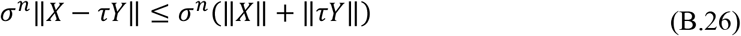

Obviously, 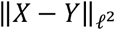 is bounded, and we have:

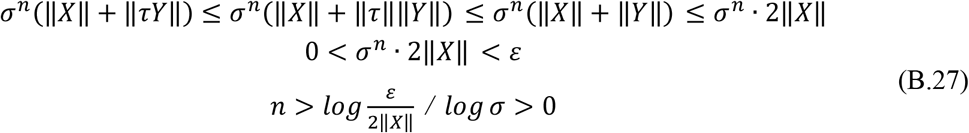

We provide the infimum of iteration as 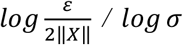 to approach the accuracy level *ɛ*.

### Theorem 2.8 (Operator PCA is bounded)

If we denote the updating operator as 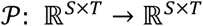, we have 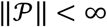.

***Proof***: According to the definition of operator norm (Rudin, 1973), 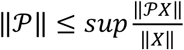; obviously, 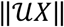 and ‖*X*‖ is bounded, since both of norms are based on finite dimensional matrix. And if we denote:

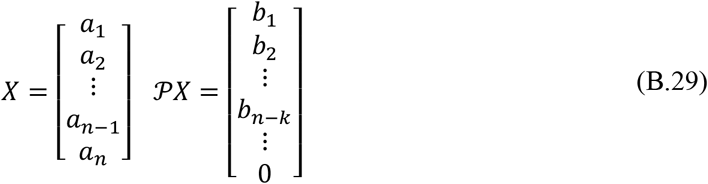

According to the dimensional reduction of PCA, we have: 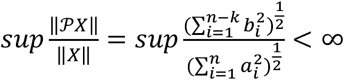. It demonstrates: 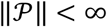.

### Theorem 2.9 (Contraction of Fixed-Point Operator)

If we denote the updating operator as 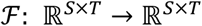, we have 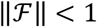.

***Proof***: according to the definition of operator norm (Rudin, 1973), 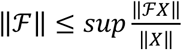; obviously, 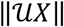 and ‖*X*‖ is bounded, since both of norms are based on finite dimensional matrix. And if we denote:

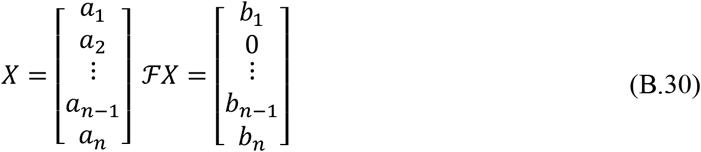

According to the iterative format of Deep NMF, we need to notice: 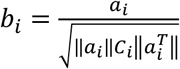; obviously, 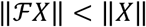. Finally, we have: 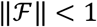. Otherwise, if 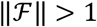, when *k* → ∞, we have: 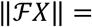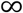.

### Theorem 2.10 (Inequality of Operator Norms)

According to Theorem 2.1, 2.7, 2.8 and 3.0, if we assume: 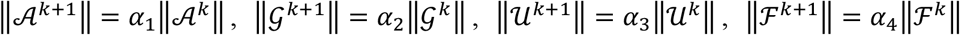, we have: α_1_ ≠ α_3_, α_4_; α_2_ ≠ α_3_, α_4_;

***Proof***: Proof by contradiction. In general, we assume 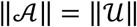, according to the iterative formats of Deep MF and Deep NMF, and considering:

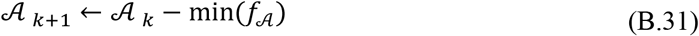

If we employ the 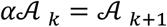 to replace 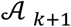:

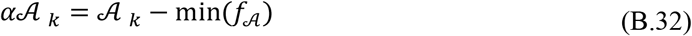

And we can reformat this equality as:

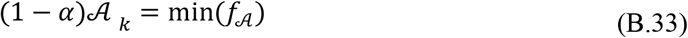

Considering the iterative format of Deep NMF:

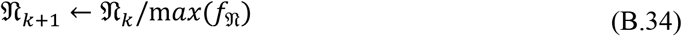

Let we denote:

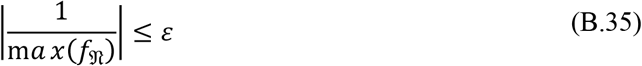

And considering an extreme condition, 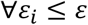, for each iteration *i*, and 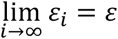;

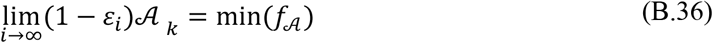

Then we have the conclusion:

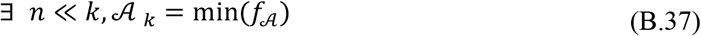

It demonstrates for the iterative format of Deep MF, before convergence, the iteration can be terminated, since a very small norm of operator 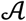.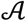 cannot guarantee the convergence. It obviously disobeys the property of ADMM.

Similarly, we can also prove α_2_ ≠ α_3_, α_4_; and α_1_ ≠ α_4_.

## Appendix C

***Assumption 3.1*** For all operators, these operators should be considered as linear operators, and we have:

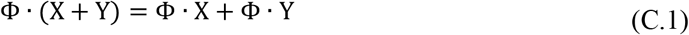

***Assumption 3.2*** For any input matrix, we can successfully separate the vital information and background noise. If we denote: 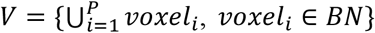, and 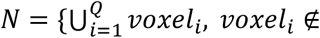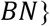. BN represents the functional areas, i.e., potentially activated areas of brain. We have some crucial assumptions: *V* ∩ *N* = ∅, *V* ≽ 0, *B* ≽ 0, ‖*V*‖ ≫ ‖*N*‖.

### Lemma 3.1 (Continuous Operators)

For all operators analyzed in this study, if *k* > *K*, ∀*k* ∈ ℕ, these iterative operators can be considered as consistent operator. It means: if we have 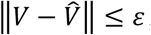,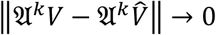.

***Proof***: We denote: 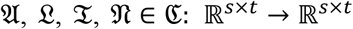

For *V*, 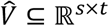, we assume that:

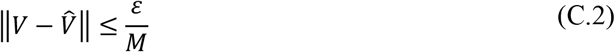

If *k* > *K*, For any operator belongs to ℭ can be considered as a contraction operator, and we have:

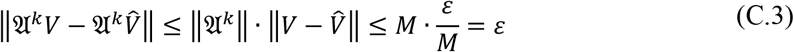

This inequality demonstrates that all operators of ℭ, if *k* is large enough, can be treated as the consistent operators (Rudin, 1973). Similarly, it also demonstrates: 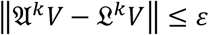

### Theorem 3.1 (Distinctive Spatial Similarity)

If we denote the following set:

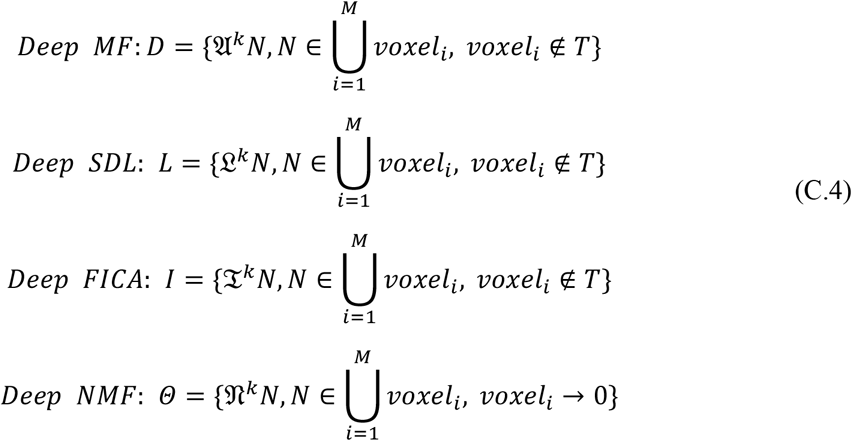

And considering the iteration *k*, it implies:

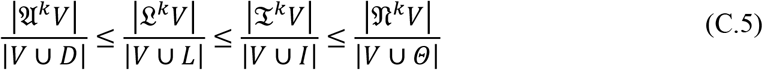

where |∙| denotes the number of positive elements.

***Proof***: Based on assumptions 3.1 and 3.2, if ∀ *k* ∈ ℕ, we have:

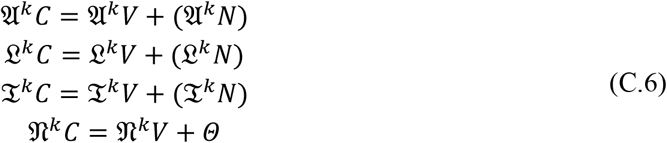

According to Corollary 1.1 and 1.2, *k* > *K*, we have:

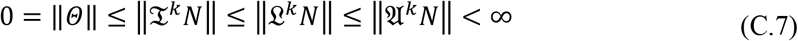

We can also rewrite it as:

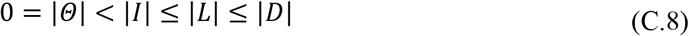

And, according to the spatial similarity, we also have:

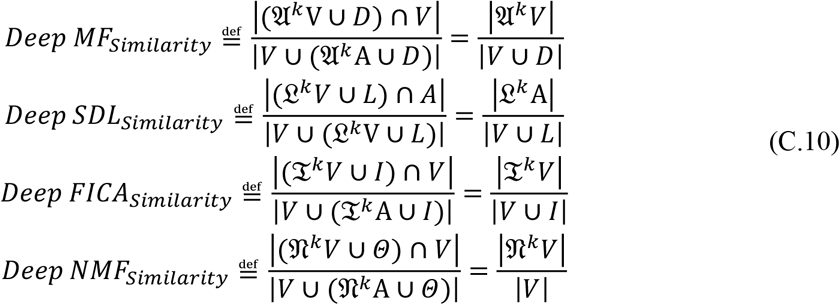

Again, considering *k* > *K*, and Corollary 1.1 to 1.2, and Theorem 3.2, we have:

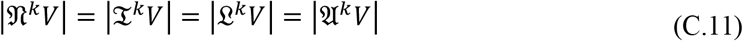

Obviously, we have:

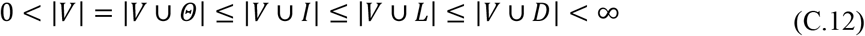

Finally, the following inequality holds, such that:

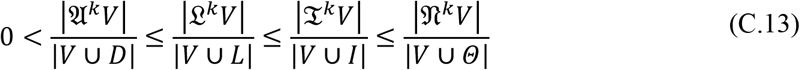

### Theorem 3.2 (Bounded Iterative Operators)

For all operators analyzed in this study, if *k* > *K*, ∀*k* ∈ ℕ, these iterative operators can be considered as consistent operator. If we have: 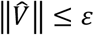, it means: 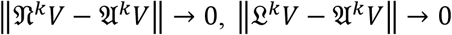 and 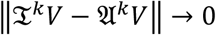.

***Proof***: We denote: 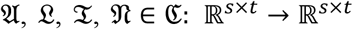

If *k* > *K*, For any operator belongs to ℭ can be considered as a contraction operator, according to Lemma 3.1, and we have:

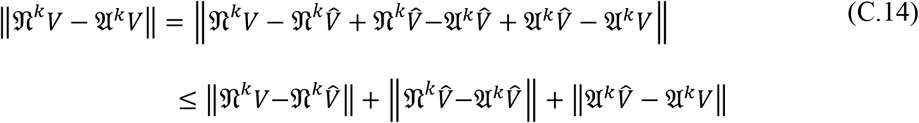

According to Lemma 3.1, and we have:

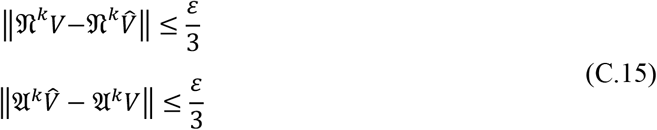

Considering the inequality:

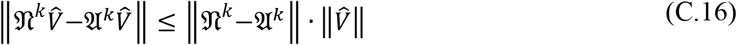

Obviously, 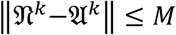, and we choose 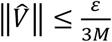; it implies:

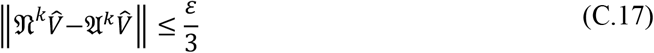

And we have:

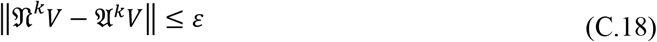

