## Supplemental Material for "Deep Linear Modeling of Hierarchical Functional Connectivity in the Human Brain"

### Supplemental Materials

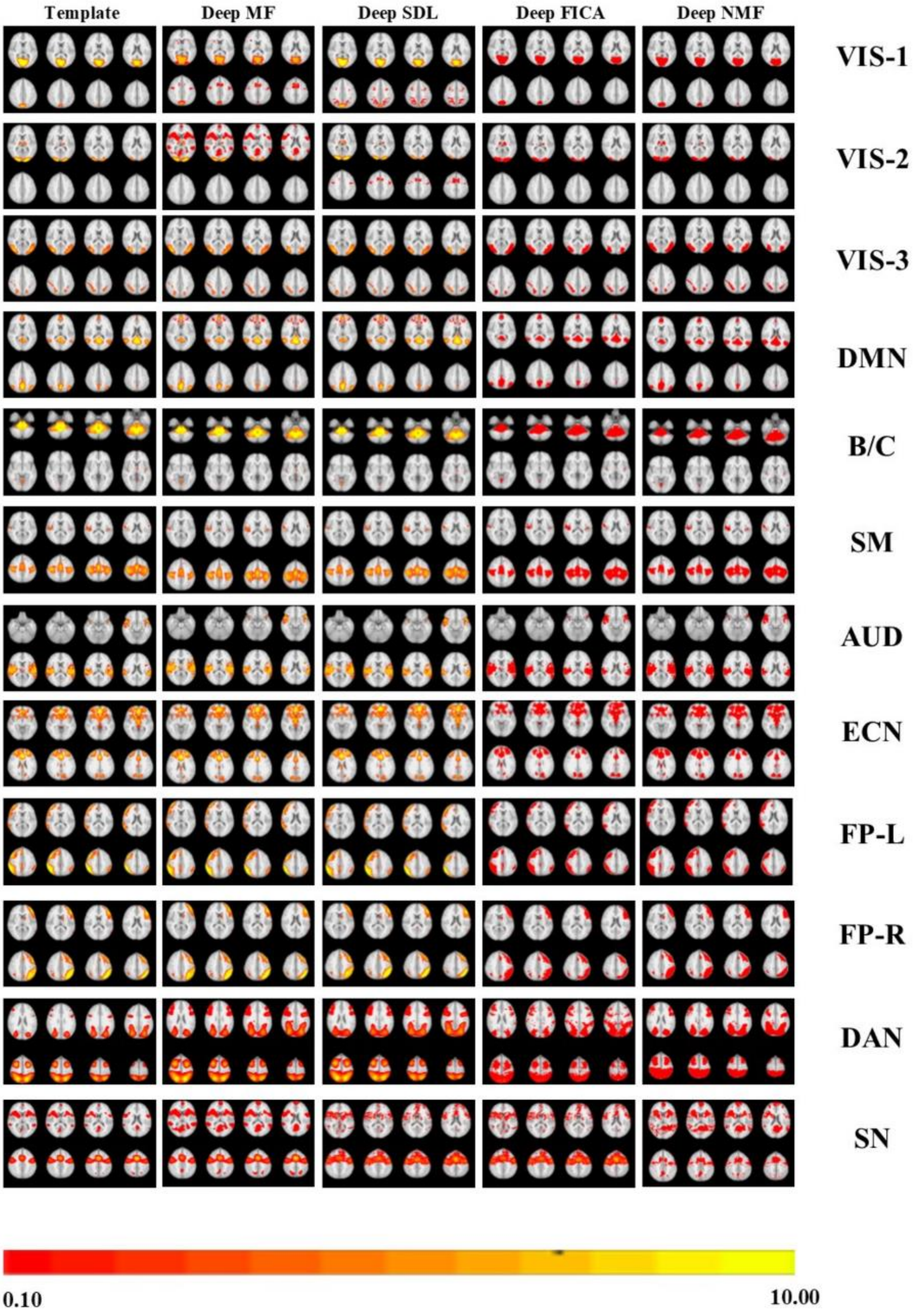

**Figure S1.** All 12 reconstructed 1st layer brain connectivity networks via Deep MF, Deep SDL, Deep FICA and Deep NMF are presented as a qualitative comparison using the most representative slices. The color bar is also provided at the bottom of this figure. The abbreviations of networks can be viewed at the right side. Each row includes the reconstructed results of a single network. The first column contains the corresponding slices of the ground truth templates. Other columns include the reconstructed results using a single deep linear model.

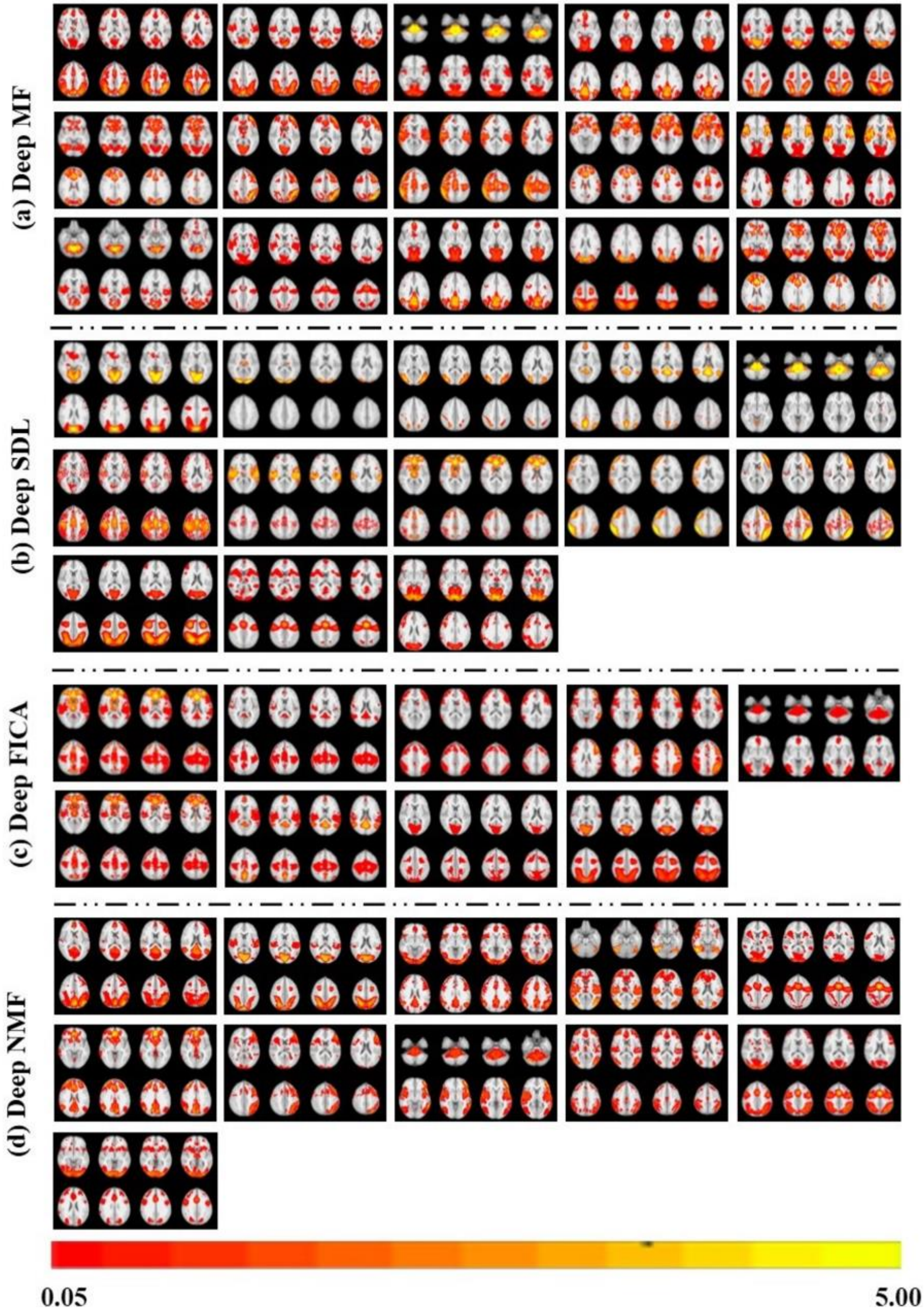

(a) Deep MF
(b) Deep SDL
(c) Deep FICA
(d) Deep NMF
0.05
5.00

**Figure S2.** All reconstructed non-noise 2nd layer BCNs via Deep MF, Deep SDL, Deep FICA and Deep NMF are presented in (a), (b), (c) and (d), respectively. The number of identified 2nd layer BCNs varies: 15, 13, 9 and 11 high-level BCNs for Deep MF, Deep SDL, Deep FICA and Deep NMF, respectively. In general, all higher-level BCNs are recombinations of shallow 1st layer features.

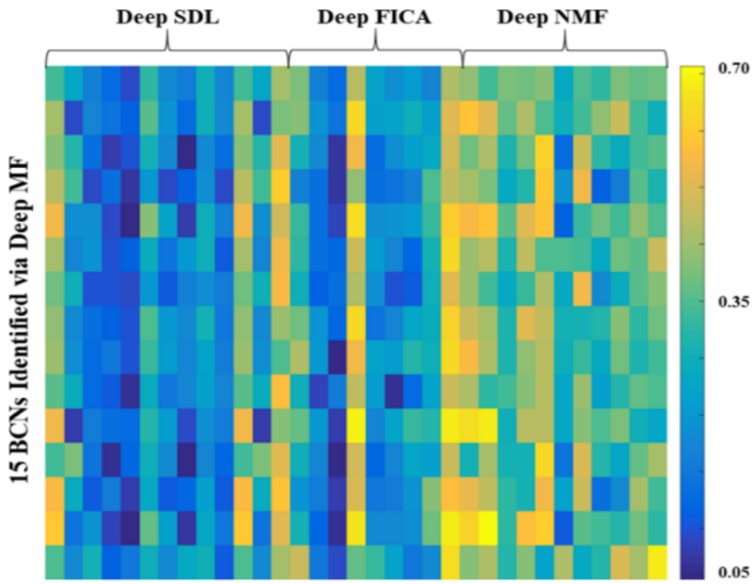


**Figure S3.** Spatial similarity matrix of all identified high-level (2$^{nd}$ layer) BCNs from 4 deep linear models. The spatial similarity of identified 2$^{nd}$ layer components recognized by Deep SDL, Deep FICA and Deep SDL (columns) versus those of Deep MF (rows) is provided as each element of the matrix. This matrix demonstrates that there are less similar high-level components identified by Deep SDL compared with Deep MF, since most of the components from Deep SDL are inherited from the 1$^{st}$ layer. However, some 2$^{nd}$ layer components recognized by Deep FICA and especially Deep NMF are more spatially similar to Deep MF, since these two deep linear models also recombine the 1$^{st}$ layer BCNs.
